# Geometric variation of the human tibia-fibula: A public dataset of tibia-fibula surface meshes and statistical shape model

**DOI:** 10.1101/2022.08.04.502722

**Authors:** Meghan Keast, Jason Bonacci, Aaron Fox

**Author notes:** Corresponding Author: Meghan Keast, Centre for Sport Research, School of Exercise and Nutrition Sciences, Deakin University 75 Pigdons Road, Waurn Ponds, 3216, VIC, Australia.

## Abstract

**Background:** Variation in tibia geometry is a risk factor for tibial stress fractures. Geometric variability in bones is often quantified using statistical shape modelling. Statistical shape models (SSM) offer a method to assess three-dimensional variation of structures and identify the source of variation. Although SSM have been used widely to assess long bones, there is limited open-source datasets of this kind. Overall, the creation of SSM can be an expensive process, that requires advanced skills. A publicly available tibia shape model would be beneficial as it enables researchers to improve skills. An opensource SSM could benefit health, sport and medicine with the potential to assess geometries suitable for medical equipment, and aid in clinical diagnosis. This study aimed to: (i) quantify tibial geometry using a SSM; and (ii) provide the SSM and associated code as an open-source dataset.

**Methods:** Lower limb computed tomography (CT) scans from the right tibia-fibula of 30 cadavers (male n = 20, female n =10) were obtained from the New Mexico Decedent Image Database. Tibias were segmented and reconstructed from the CT scans into both cortical and trabecular sections. Fibulas were segmented as a singular surface. The segmented bones (cortical and trabecular tibia; and fibula) were used to develop three SSM of the: (i) tibia; (ii) combined tibia and fibula; and (iii) tibial trabecular. Principal component analysis (PCA) was applied to obtain the three SSM, with the principal components (PCs) that explained 95% of the geometric variation retained.

**Results:** Overall size variation was the main source of variation of all three models accounting for 90.31%, 84.24% and 85.06%, respectively. Other sources of geometric variation in the tibia surface models included overall (PC3) and midshaft thickness (PC2); prominence and size of the condyle plateau, tibial tuberosity, and anterior crest (PC4); and axial torsion of the tibial shaft (PC5). Further variations in the tibia-fibula model included midshaft thickness of the fibula (PC2); fibula head position relative to the tibia (PC2); tibia and fibula anterior-posterior curvature (PC4); fibula posterior curvature (PC5); tibia plateau rotation (PC5); and interosseous width (PC6). The main sources of variation in the tibia-trabecular model other than general size included variation in the medulla cavity diameter (PC2); overall thickness (PC3); and the relative volume of proximal and distal ends compared to middle (PC4).

**Conclusion:** Important variations that could increase the risk of tibial stress injury were observed in the tibia and tibia-fibula SSM. These included general tibial thickness, midshaft thickness, tibial length and medulla cavity diameter (indicative of cortical thickness). Further research is warranted to better understand the effect of these tibial and fibula shape characteristics on tibial stress, loading and injury risk. This SSM and associated code has been provided in an open-source dataset. The associated code includes three example applications: (i) generation of a random sample; (ii) reconstruction of trabecular surfaces; and (iii) reconstruction from palpable landmarks. The developed tibial surface models and statistical shape model will be made available for use at: https://simtk.org/projects/ssm_tibia.

## Introduction

Statistical shape models (SSM) offer a method to assess three-dimensional variation of structures, including the sources of variation in bones. Although SSM have been used widely to assess long bones (Zhang et al., 2014a; Nolte et al., 2020; Bruce et al., 2021) there is limited open-source datasets of this kind. SSM that use medical imaging can be costly and the funding or facilities needed to produce high quality imagery can be inaccessible. Segmentation of medical images and bone surface reconstruction requires specialized software, skill, and extensive time. Further, transforming the segmented data into three-dimensional models requires an intermediate level of computer coding knowledge. Attainment of coding proficiency is a steep learning curve that requires extensive mentorship. Overall, the creation of SSM can be an expensive and time costly process, that requires advanced skills. A publicly available tibial shape model would be beneficial as it enables researchers who may be affected by the aforementioned hurdles to improve skills and adapt the code to meet their needs. Openly sharing code, data and instructional papers is beneficial to all fields of research (Pisani et al., 2016; Pronk, 2019). SSM can also further research in health, sport, and medicine with the potential to create a sample of computer simulated tibias, assess geometries suitable for medical equipment, and aid in clinical diagnosis.

The ability to characterize and understand bone shape and geometric variation is also an important aspect in skeletal research. Bone stress injuries caused by exercise are one area that may benefit from an increased understanding of bone shape and individual variation. Individuals who engage in high volumes of running, through recreational running or sport participation, are at an elevated risk for lower limb bone stress injuries (Rizzone et al., 2017). Tibial stress reactions and tibial stress fractures are the fifth and ninth most common running-related injuries, respectively (Taunton, 2002). Tibial stress injuries result, in part, from the mechanical fatigue of bone after cyclic loading (Burr et al., 1991). Tibial stress injuries are multifactorial in nature and can be caused by several intrinsic and extrinsic factors, one of these being skeletal geometry (Popp et al., 2019). Variations in tibial geometry such as smaller tibias (Crossley et al., 1999; Beck et al., 2014) and a thinner mid diaphysis (Popp et al., 2019) have been cited as a risk factor for tibial stress injury. Quantifying tibial anatomy using SSM and providing this as an open resource can assist in progressing our understanding of risk factors for tibial stress injury, as well as identify relevant interventions to reduce the risk of these injuries in running populations. Therefore, the aims of this paper are: (i) to quantify tibial geometry using a SSM; and (ii) provide the SSM and associated code as an open-source dataset. This paper and associated code will also demonstrate potential applications for the SSM to guide potential use.

## Methods

Lower limb computed tomography (CT) scans from the right tibia and fibula of 30 cadavers (male n = 20, female n =10) were obtained from the New Mexico Decedent Image Database (Edgar et al., 2020). The images included were from individuals with a mean (± standard deviation) age of 28.7± 6.7, living weight of 70.22 ± 11.36kg, and living height of 176.06 ± 11.61cm. Individuals whose records indicated participation in impact-based physical activities throughout life were selected for inclusion (i.e. team sports, dancing, recreational running and walking). This criterion was included in an effort to ensure participants were sampled from a generally active population. Participants were excluded if there was any noticeable damage to the right tibia-fibula or the inclusion of medical devices (e.g. plates, rods, or screws). The tibia and fibulas were segmented from the CT images using Mimics innovation suite (Materialise, Leuven, Belgium). Tibias were segmented into two surfaces representing the outer boundaries of the trabecular and cortical bone, while fibulas were segmented as one surface representing the outer shape of the entire bone. Cortical bone was classified as all visible hard bone, while trabecular was all remaining soft bone and the medulla cavity. The medulla cavity was included in the trabecular bone segmentation as this enabled us to describe changes in size and shape of the entire trabecular surface and medulla cavity without needing to create a cross section of the tibia. Segmentation was done using a combination of image intensity thresholding with manual corrections. Landmarks on the lateral and medial malleoli, lateral and medial tibial condyle, fibula head, tibial tuberosity, anterior aspect of the tibia at 25%, 50% and 75% of the distance between the medial condyle and malleolus, and lateral fibula diaphysis 25% proximally from the malleoli were registered from the medical images on the segmented surfaces (Bruce et al., 2021).

The segmented bones (cortical and trabecular tibia; and fibula) were used to develop three SSM of the: (i) tibia; (ii) combined tibia and fibula; and (iii) tibial trabecular. The procedures used to create the SSM were applied identically across all three models. Prior to development of the SSM, the bones were aligned to the tibial reference system as per International Society of Biomechanics recommendations (Wu et al., 2002). This was done to simplify later steps in the surface registration processes. Surfaces were remeshed to have a matching number of points (tibia *n* = 3500; fibula *n* = 1500; trabecular *n* = 1500). Nodal correspondence and registration were performed in MATLAB (R2019b, Mathworks, MA, United States). All surfaces were registered against the same target mesh, which was selected due to being the closest to the sample mean (Bruce et al., 2021). Nodal correspondence was performed using the coherent drift point algorithm (Myronenko & Song, 2010). This algorithm non-rigidly registers (i.e. rotates, translates and scales) each surface mesh to the target mesh so that corresponding points between surfaces can be identified. The surface meshes were then rigidly aligned using a Procrustes analysis with no scaling applied.

Principal component analysis (PCA) was applied to the surface sets to obtain the three SSM. The principal components (PCs) that accounted for > 95% of geometric variation in each SSM were retained (Smoger et al., 2017). Mean and peak error in surface node position, and the Jaccard index (i.e. a measure of volumetric similarity where values range from 0 to 1 indicating no to perfect similarity, respectively) (Real & Vargas, 1996)(mean ± 95% confidence intervals [CIs]) of reconstructed surfaces using the reduced component set were calculated for each SSM. To interpret the variation explained by each PC, the mean surface was compared to surfaces generated by manipulating the retained PC up to plus/minus three standard deviations. Animated heatmaps of each PC component, where colour variation indicated the relative amount of position change of the surface nodes, were also used to assist with interpretation of PCs (see supplementary files 1, 2 and 3).

## Results

### Tibia SSM

The first five PCs accounted for 95.80% of the geometric variation across the tibial surfaces (see Table 1). Mean and peak error in surface node position (in mm), and the Jaccard index for the reconstructed tibia surfaces using a reduced component set were 1.76 [1.66, 1.86 95% CIs], 4.70 [4.41, 4.98 95% CIs] and 0.773 [0.771, 0.776 95% CIs], respectively (see Figure 1).

**Table 1.**
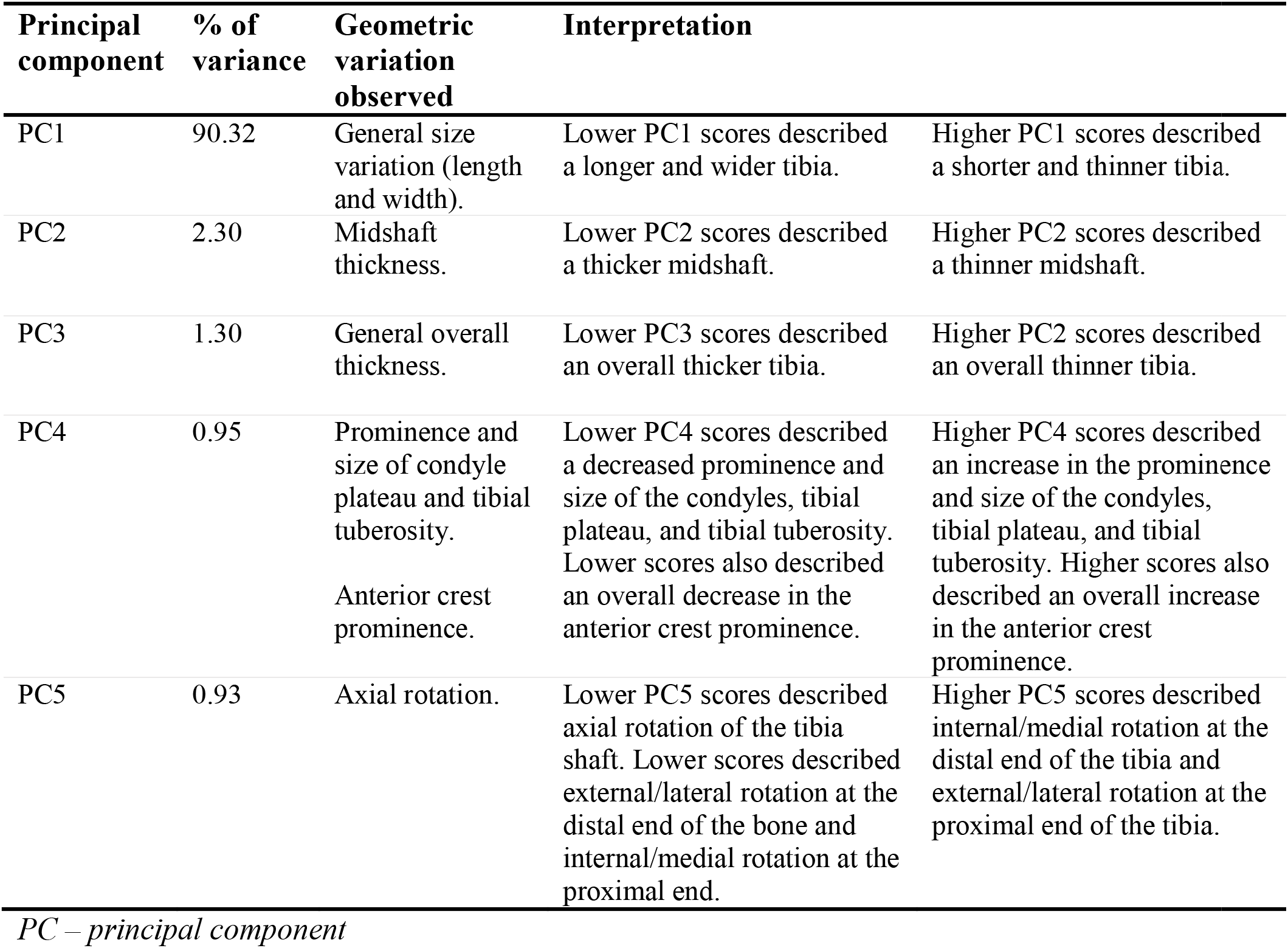
Percentage of variance and interpretations of each principal component for the tibia SSM.

**Figure 1.**
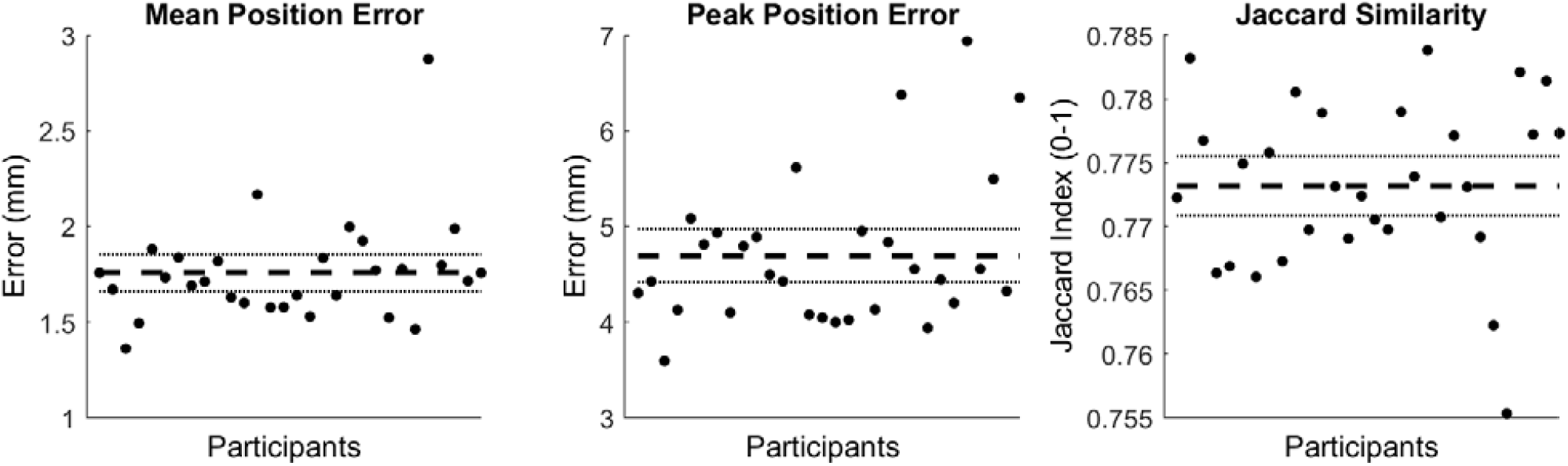
Mean and peak error in surface node position, and the Jaccard index for the reconstructed tibia surfaces using a reduced component set. Error variables are presented in mean (dashed line) and 95% confidence intervals (dotted lines).

### Tibia - Fibula SSM

The first seven PCs accounted for 95.49% of the geometric variation across the tibial surfaces (see Table 2). Mean and peak error in surface node position (in mm), and the Jaccard index for the reconstructed tibia-fibula surfaces using a reduced component set were 1.89 [1.81, 1.97 95% CIs], 5.67 [5.30, 5.99 95% CIs] and 0.751 [0.743, 0.759 95% CIs], respectively (see Figure 2).

**Table 2.** Percentage of variance and interpretations of each principal component for the tibia-fibula SSM.

| Principal component | % of variance | Geometric variation observed | Interpretation |  |
| --- | --- | --- | --- | --- |
| PC1 | 84.24 | General size variation (length and width) of tibia and fibula. | Lower PC1 scores described a longer and wider tibia and fibula | Higher PC1 scores described a shorter and thinner tibia and fibula. |
| PC2 | 3.50 | Midshaft thickness of fibula.<br>Fibula head position. | Lower PC2 scores described a thinner fibula midshaft, with a more anterior head relative to the tibia. | Higher PC2 scores described a thicker fibula midshaft, with a more posterior head relative to the tibia. |
| PC3 | 2.60 | General overall thickness of tibia and fibula. | Lower PC3 scores described an overall thicker tibia and fibula, | Higher PC3 scores described an overall thinner tibia and fibula. |
| PC4 | 1.91 | Tibia and Fibula anterior-posterior curvature. | Lower PC4 scores described posterior curvature of the fibula at the proximal end, the fibula head becomes more anterior. Lower scores also describe a straighter tibia. | Higher PC3 scores describe a straighter fibula, with a more posterior head. The tibia displays anterior curvature. |
| PC5 | 1.36 | Fibula posterior curvature.<br>Tibia plateau rotation. | Lower PC5 scores describe a straighter fibula, flattening in an anterior direction. Lower scores also describe internal rotation of the tibial plateau. | Higher PC5 scores describe a more posteriorly curved fibula, and a more externally rotated tibial plateau. |
| PC6 | 1.03 | Tibia upper to midshaft thickness.<br>Tibia condyle prominence. | Lower PC6 scores describe a thinner tibia, with a decreased condyle prominence. | Higher PC6 scores describe a thicker tibia with increased condyle prominence. |
| PC7 | 0.85 | Interosseous width. | Lower PC7 scores describe a smaller/thinner gap between the tibia and fibula at the distal end. | Higher PC7 scores describes a bigger/thicker gap between the tibia and fibula at the distal end. |

**Figure 2.**
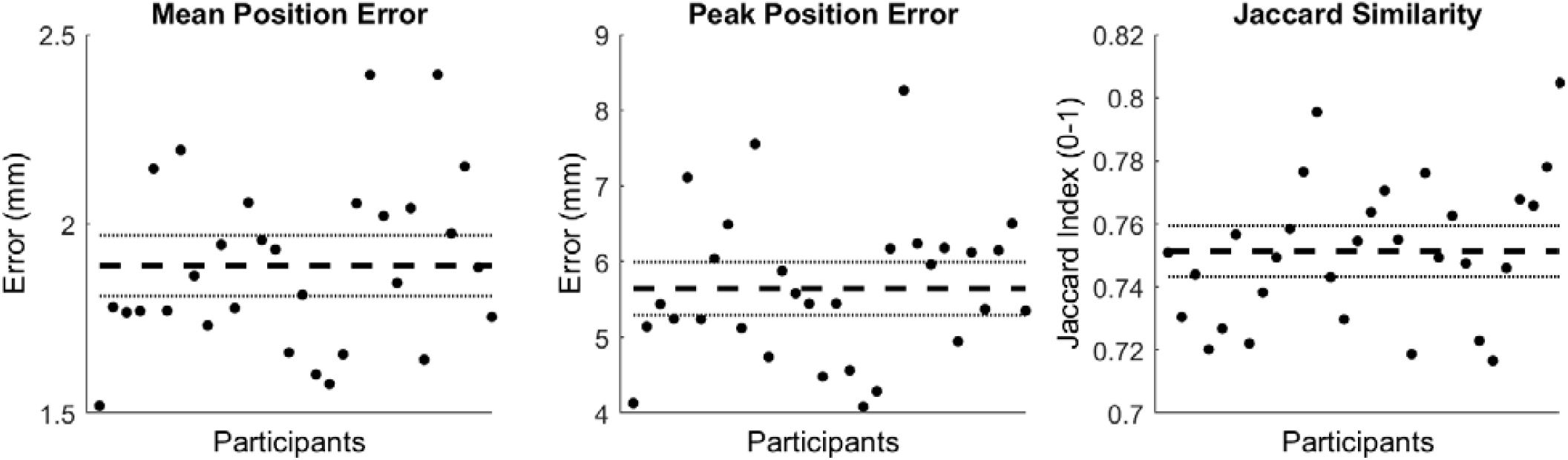
Mean and peak error in surface node position, and the Jaccard index for the reconstructed tibia-fibula surfaces using a reduced component set. Error variables are presented in mean (dashed line) and 95% confidence intervals (dotted lines).

### Tibia - Trabecular SSM

The first four PCs accounted for 95.17% of the geometric variation across the tibia trabecular (see Table 3). Mean and peak error in surface node position (in mm), and the Jaccard index for the reconstructed trabecular surfaces using a reduced component set were 2.05 [1.96, 2.14 95% CIs], 5.63 [5.31, 5.95 95% CIs] and 0.759 [0.753, 0.764 95% CIs], respectively (see Figure 3).

**Table 3.** Percentage of variance and interpretations of each principal component for the trabecular SSM.

| Principal component | % of variance | Geometric variation observed | Interpretation |  |
| --- | --- | --- | --- | --- |
| PC1 | 85.07 | General size variation (length and width). | Lower PC1 scores describe a longer and wider trabecular bone. | Higher PC1 scores describe a shorter and thinner trabecular bone. |
| PC2 | 7.50 | Medulla cavity diameter. | Lower PC2 scores describe an increased diameter of the medulla cavity. | Higher PC2 scores describe a decreased diameter of the medulla cavity. |
| PC3 | 1.50 | General overall thickness | Lower PC3 scores describe an overall thicker trabecular bone. | Higher PC3 scores describe an overall thinner trabecular bone. |
| PC4 | 1.10 | Relative volume of proximal and distal ends compared to the middle. | Lower PC4 scores describe an increased trabecular bone volume at the ends of the bone compared to the middle of the bone. This indicates a decreased medulla cavity diameter and a potential thicker cortical bone at the midshaft of the tibia. | Higher PC4 scores describe a more consistent volume across the bone, this indicates an increased medulla cavity diameter and potentially thinner cortical bone at the midshaft of the tibia. |

*PC – principal component*

**Figure 3.**
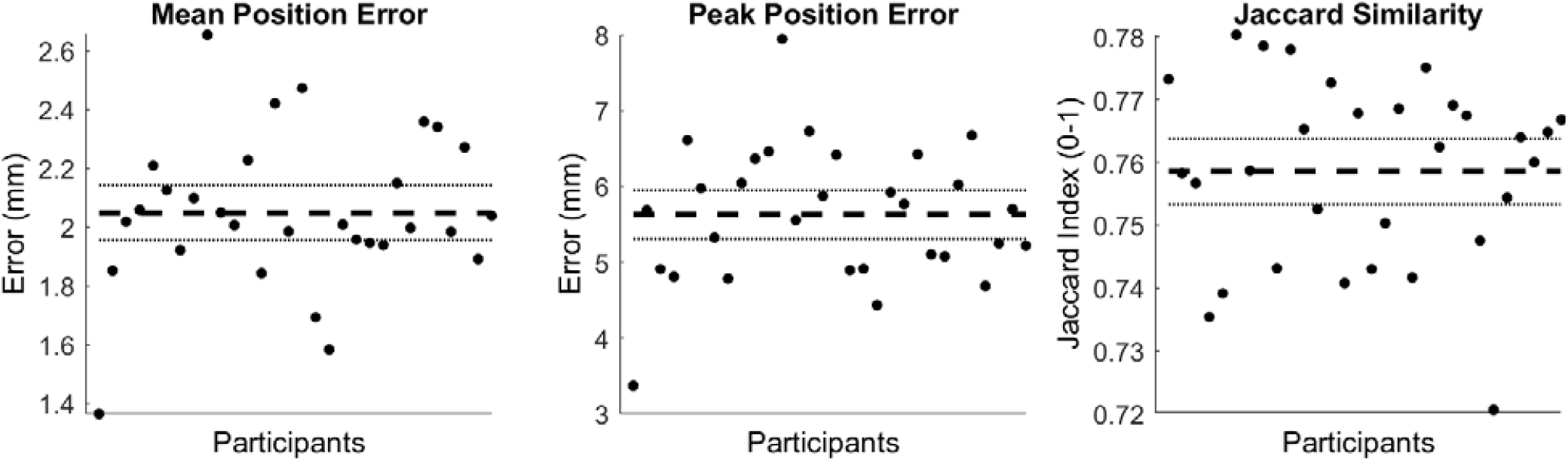
Mean and peak error in surface node position, and the Jaccard index for the reconstructed tibia-trabecular surfaces using a reduced component set. Error variables are presented in mean (dashed line) and 95% confidence intervals (dotted lines).

## Discussion

The purpose of this study was to explore tibial geometry using a SSM and provide the SSM and associated code as an open-source dataset. The data presented and associated code can be found online at https://simtk.org/projects/ssm_tibia. Our results indicated that overall size variation (length and width) was the main source of variation of all three models (i.e. tibia, tibia and fibula, and tibial trabecular) accounting for 90.31%, 84.24% and 85.06%, respectively. This aligns with previous SSM of the tibia where overall size was the main source of variation in tibia (Quintens et al., 2019) and tibia-fibula models (Bruce et al., 2021). Other sources of geometric variation in our tibia surface models included overall and midshaft thickness; prominence and size of the condyle plateau, tibial tuberosity and anterior crest; and axial torsion of the tibial shaft. Sources of variation noted in the tibia model were also evident in the tibia-fibula model, however inclusion of the fibula introduced variations in midshaft thickness of the fibula; fibula head position relative to the tibia; tibia and fibula anterior-posterior curvature; fibula posterior curvature; tibia plateau rotation; and interosseous width. However, these variations only accounted for a small proportion of overall tibia-fibula geometric variation. Lastly, sources of variation in the tibia-trabecular model included variation in the medulla cavity diameter; overall thickness; and the relative volume of proximal and distal ends compared to middle. Previous research has identified both similar and different variations in tibial geometry compared to the outcomes from our research(Tümer et al., 2019; Quintens et al., 2019). Similar to the variations identified in our tibia model, past studies identified variations in the ‘bow’ of the tibia in the anterior posterior direction (i.e. anterior posterior shaft curvature) (Quintens et al., 2019), as well as changes in the orientation of the tibial plateau (Quintens et al., 2019). Variation in the prominence of the condyles as well as changes to the tibial tuberosity and overall thickness of the tibia were also evident in past isolated studies of the tibia (Tümer et al., 2019; Quintens et al., 2019).

Several of the observed variations in tibial geometry could relate to increased risk of tibial stress injury. Smaller and thinner tibial geometry was the predominant source of variation found in our SSM, which can impact the bones’ ability to withstand bending forces created during exercise (Nordin & Frankel, 2012). The impact that a smaller tibia has on tibial stress injuries has been cited in both athletic (Crossley et al., 1999; Beck et al., 2014) and military populations (Giladi et al., 1991; Beck et al., 2000). However, size of bone is often proportionate to individuals’ height and mass (Duyar & Pelin, 2003). The population used for this study had a height and mass range of 160 to 200cm and 49.9 to 93kg, respectively. It is likely that the smaller tibias we observed were related to height and mass and this may not automatically be indicative of elevated tibial stress injury risk. Despite having smaller tibias, shorter individuals would likely produce reduced forces and moments during running given their lighter mass and bone segment length. Nonetheless, our shape model demonstrates that tibial size, a commonly cited risk factor for tibial stress injuries (Giladi et al., 1991; Crossley et al., 1999; Beck et al., 2000, 2014), is a primary source of variation in tibial geometry. What is likely problematic is when an individual with greater height/mass demonstrates these smaller tibial properties. Reducing tibial forces and moments during running in individuals displaying this anatomical characteristic is likely beneficial for minimising tibial stress injury risk.

PC2 of the trabecular model described the diameter of the medulla cavity. Changes in the diameter could indicate changes in overall bone width in addition to thickness of the cortical bone (i.e. enlarged medulla cavity equating to reduced cortical bone thickness). Cortical bone thickness is cited as a risk factor for tibial stress injuries in military (Cosman et al., 2013) and athletic populations (Beck et al., 2014). Similar to smaller and thinner tibial geometries, decreased cortical thickness could impact a bones ability to withstand loads. Reduced thickness of the cortical bone could reduce the overall stiffness of the tibia and make it more susceptible to compression and bending forces during exercise (Nordin & Frankel, 2012). PC3 of both the tibia and tibia-fibula model described changes in overall thickness of the tibia, while PC2 of the tibia model described changes in midshaft thickness. General tibia thickness may be an important factor for tibial stress injuries (Giladi et al., 1991). However, tibial midshaft thickness is likely a more important variation than overall tibia thickness for tibial stress injury risk. Tibial stress injuries most commonly occur in the mid to distal third of the tibial shaft (Coady & Micheli, 1997; Beck, 1998). A past study found that when adjusting for body size, thinner mid–diaphysis of the tibia is an important factor in the incidence of tibial stress injuries (Popp et al., 2019). Decreased overall tibial thickness and thickness of the midshaft likely decrease the tibias’ ability to withstand bending forces created during exercise, potentially increasing the risk of tibial stress injury. Promoting running technique that minimizes tibial forces and moments in individuals with these geometries could likely be important for reducing tibial stress injury risk.

Additional areas of geometric variation we observed in the tibia, such as axial torsion of the tibia, interosseous width, and anterior-posterior curvature have not been considered in relation to tibial stress injury risk. Further, we have limited understanding of how changes in fibula geometry influence tibial stress. Previous research has identified that the fibula potentially acts in a bracing fashion, by restricting medial and posterior motion of the tibial plateau relative to the malleoli (Haider, Baggaley & Edwards, 2020). It is unknown if changes in fibula geometry might impact its ability to provide this support. Further research is warranted to better understand the effect of these additional tibial shape characteristics and fibula geometry on tibial stress, loading and injury risk.

### Practical Applications of the SSM

The dataset and associated code provided at https://simtk.org/projects/ssm_tibia can be used in several ways to assist with skeletal focused research of the tibia-fibula complex. This paper does not have the scope to highlight each and every potential application, however we propose three broad applications of where and how the SSM may be used. The code and analyses described in the following case studies are also included with our dataset.

### Case One: Generating Surface Samples

Obtaining CT and/or MRI scans to develop a large sample of bone surfaces can be costly from both a time and financial perspective. Further, the imaging facilities to obtain these data may not be available to all. In this case study, we provide an application that uses the SSM of the tibia-fibula to create a sample set of tibia-fibula surfaces representing standardized variation across all and/or a selection of the model components. The ‘simulatedPopulations.m’ function provided with our dataset allows users to generate a select number of surfaces from a pre-loaded shape model. While the example provided uses the tibia-fibula shape model, this process could be applied to any of the shape models included in our dataset. The function is structured to allow users to: (i) load the desired shape model; (ii) set the desired number of samples (i.e. *n*) to be generated; (iii) select the components of the model to be perturbed in creating the surface variations; (iv) set the magnitude of variation to perturb the model components by (in standard deviations); and (v) set the output options. The function takes these user inputs and randomly samples values for the chosen components within the standard deviation bounds provided to reconstruct *n* samples. Users can select to output a visual representation of the surfaces with heatmaps applied to highlight the areas of variation (see Figure 4), as well as outputting the three-dimensional surfaces in STL format.

**Figure 4.**
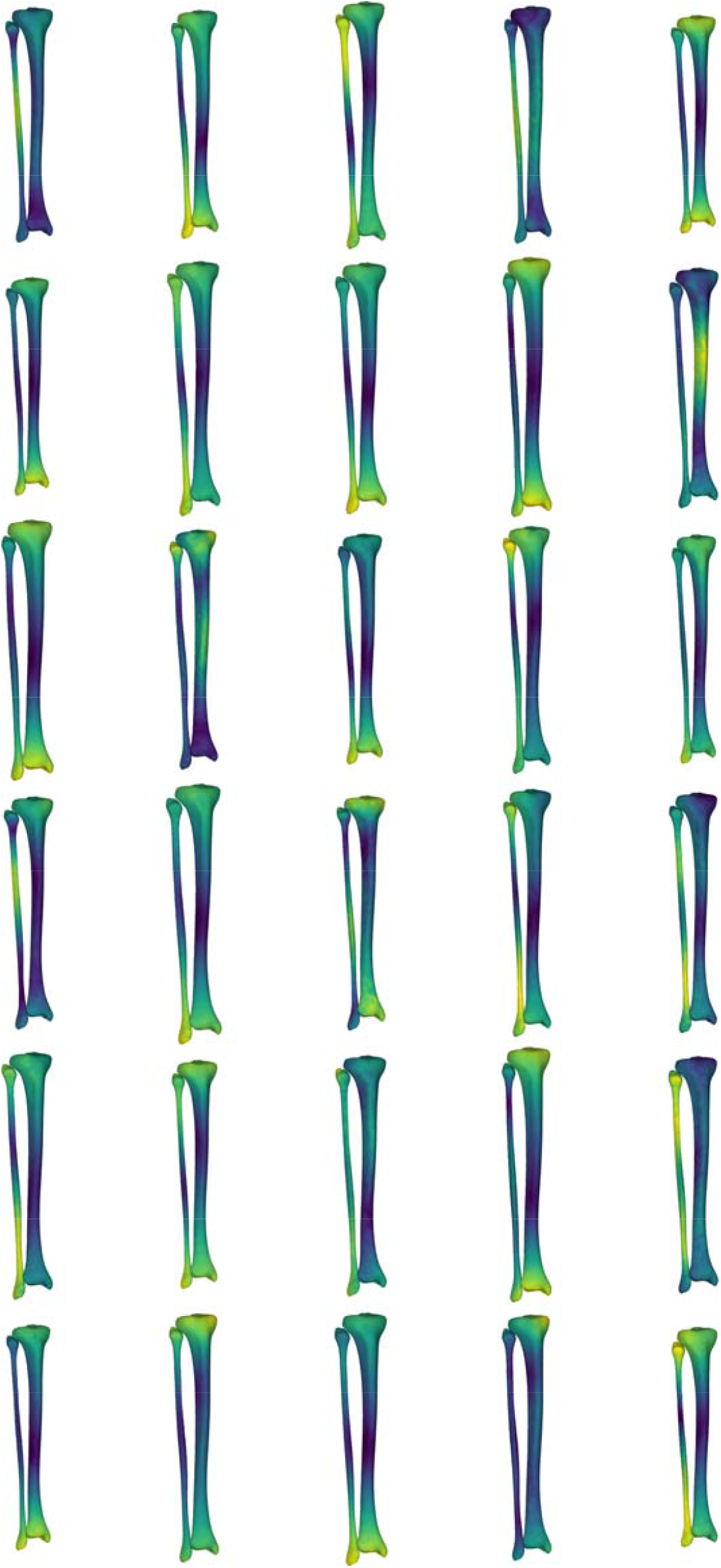
Simulated surface sample of 30 tibia-fibulas, this sample was created using the first five and the seventh principal components, opting to ignore the sixth. Standard deviation bounds were set at 1.5. Each surface is displayed with a corresponding heatmap, detailing the changes that occurred with each sample. Warmer colours describe greater change occurring at that area of tibia-fibula, respective to the mean shape model.

The SSM developed in our study represent the major sources of variation in the bone surfaces, and this model can be used to produce a representative sample of surfaces or a ‘simulated population.’ The ability to perturb or manipulate specific components of the shape models also allows for samples to be generated with isolated geometric variations of interest (e.g. specifically isolating variation in mid-shaft tibial thickness).

### Case Two: Predicting and Generating Trabecular Volumes

Finite element simulations of tibial loading typically include different parts and material properties for the cortical and trabecular volumes of the bone (Edwards et al., 2010; Haider, Baggaley & Edwards, 2020). Certain scenarios can make generating surface and volumetric meshes of the trabecular more difficult or time consuming. Segmenting the trabecular surface from CT scans is possible, yet more time-consuming given the need for greater manual corrections. MRIs that are not optimized to detect the trabecular volume can also generate difficulties in segmenting the trabecular area due to lacking clearly identifiable borders between the cortical and trabecular areas. Other bone surface reconstruction methods (e.g. MAP-client (Zhang et al., 2014b)) only provide estimates of external bone surface and not internal trabecular volumes. Given that performing finite element simulations with both trabecular and cortical bone is more reflective of real-world scenarios, there are practical benefits to developing methods that can predict and generate trabecular volumes based on the more easily obtainable outer bone surface.

In this case study, we use the SSM of the tibia and trabecular to develop and evaluate a method for estimating a correspondent trabecular surface. The theoretical hypothesis for this approach is that the outer shape of the tibia can be effective in predicting the internal trabecular shape. The ‘generateTrabecularFromSurface.m’ function provided with our dataset demonstrates the development and evaluation of this method on the dataset used in the present study, followed by the application of the method to a new set of tibial surfaces. Briefly, this code develops a set of linear regression models (see supplementary file 4) using the retained principal component scores from the tibia model to predict each of the retained principal component scores of the trabecular model. The predicted scores can then be combined with the trabecular SSM to reconstruct the trabecular surface. In developing this method we noted that it was common for the reconstructed trabecular to project outside of the tibial surface (i.e. not fitting within the outer tibia boundary). To combat this, we included a final step where points on the reconstructed trabecular surface that protruded outside the tibia surface were translated to the closest point one millimetre inside the boundary of the tibia.

We evaluated the accuracy of the trabecular reconstruction method on the surfaces used to create the SSM using a leave-one-out approach. Specifically, the linear regression models were iteratively developed while leaving out one of the participants surfaces – and the reconstruction accuracy of the ‘left-out’ trabecular surface evaluated using the mean and peak error in surface node position, alongside the Jaccard index. Mean and peak error in surface node position (in mm), and the Jaccard index for the reconstructed trabecular surfaces using the prediction model were 3.05 [2.64, 3.47 95% CIs], 8.38 [7.21, 9.54 95% CIs] and 0.756 [0.750, 0.762 95% CIs], respectively (see Figure 5). Figure 7 presents an ‘average’ performing reconstruction (i.e. surface with Jaccard index closest to the mean) of the trabecular surface compared to the originally segmented trabecular for this participant.

**Figure 5.**
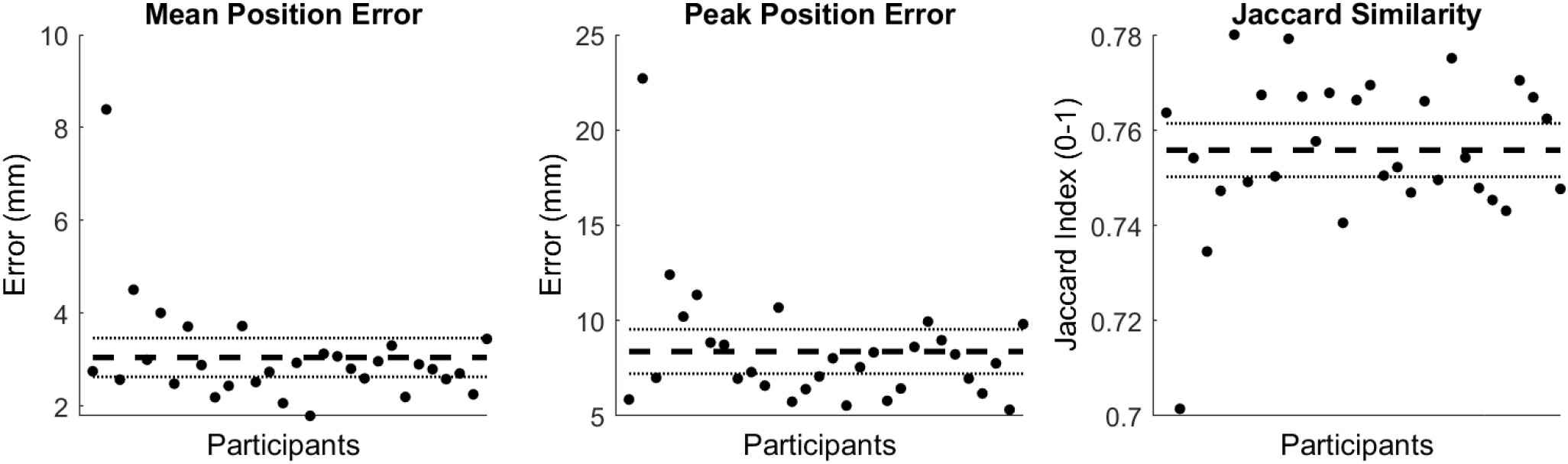
Mean and peak error in surface node position, and the Jaccard index for the reconstructed trabecular surfaces using the prediction model. Error variables are presented in mean (dashed line) and 95% confidence intervals (dotted lines).

**Figure 6.**
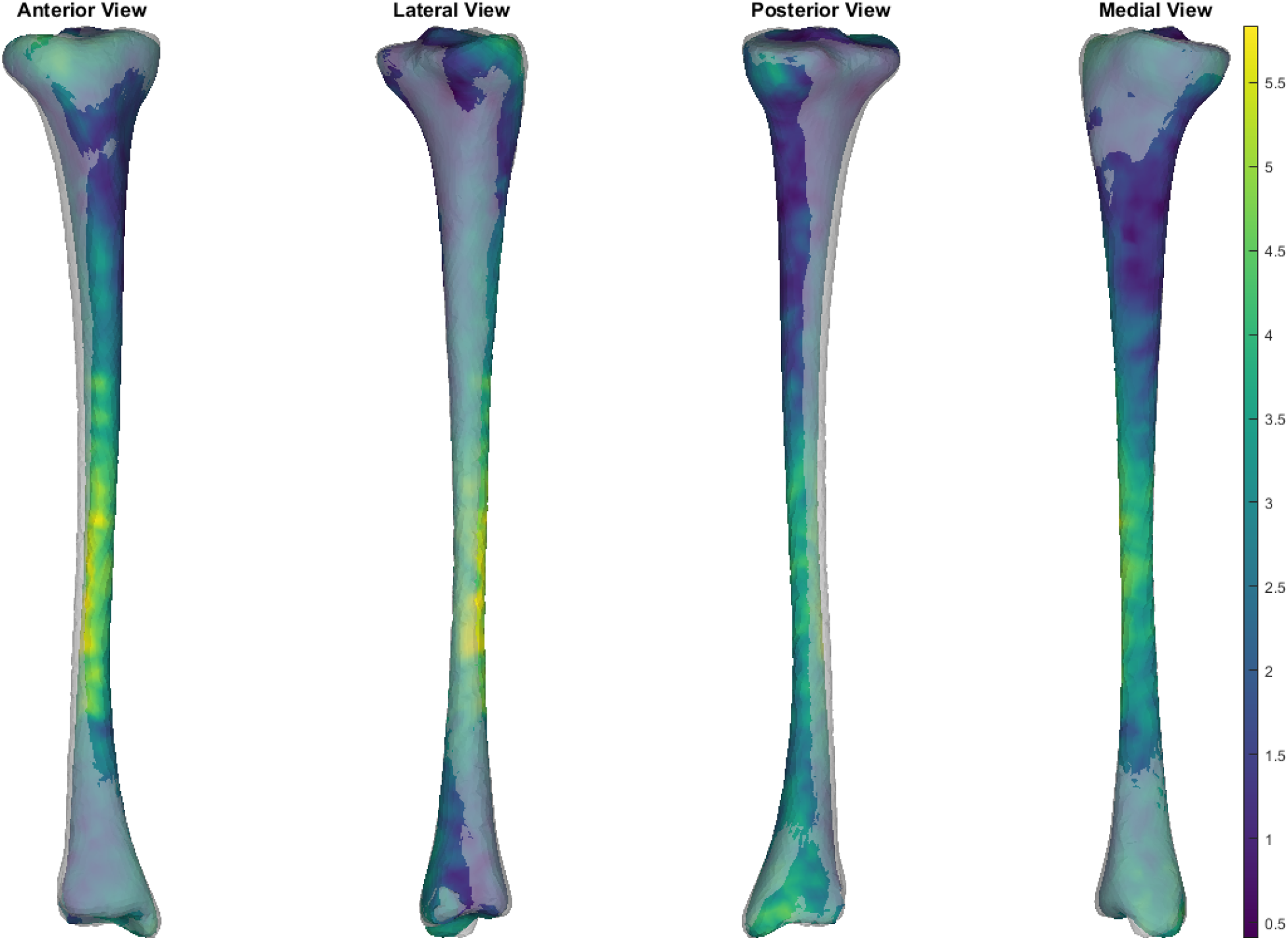
Comparison of a reconstructed trabecular surface (i.e. surface with Jaccard index closest to the mean) compared to the originally segmented trabecular for this participant. The transparent grey surface is the original segmentation and the colour map is the predicted trabecular surface. Warmer colours represent greater point distance (mm) from the original segmentation.

**Figure 7.**
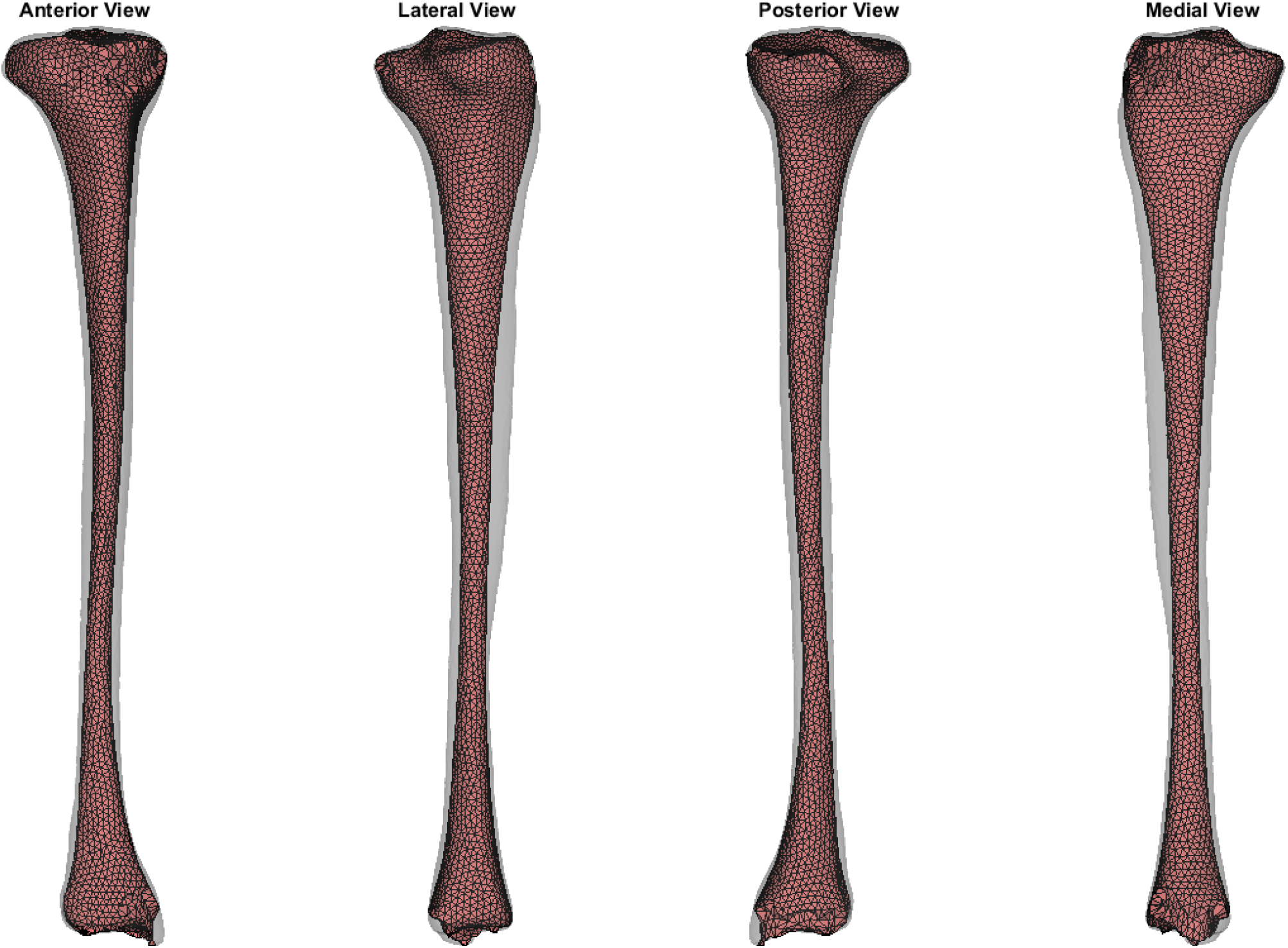
Predicted trabecular surface (red) of a randomly selected tibia (transparent grey) from Notle et al., (2016).

This case study also provides an example of how this method can be applied to estimate trabecular volumes on newly acquired tibial surface (Nolte et al., 2016). The trabecular of a randomly selected sample from the tibia-fibula surfaces provided by Nolte et al., (2016) was predicted using the shape models from the present study (see Figure 7). The relative accuracy of these specific trabecular reconstructions cannot be evaluated due to Nolte et al., (2016) only providing the outer tibial surface meshes. Qualitative evaluation of the predicted trabecular surface from this new sample revealed potential quality issues towards the ends of the bone. Nonetheless, the code is provided to demonstrate how this method can be applied to any tibial surface data with the caveat that quality control checks following predictions are likely required.

### Case Three: Generating Tibia-Fibula Surfaces from Palpable Landmarks

Providing methods to generate three-dimensional models of the tibia-fibula allow those who cannot acquire medical images to use these models within their work or practice. Obtaining spatial measurements from palpable landmarks on the tibia-fibula surface is a much more accessible method and can be achieved through motion capture or video-based methods. Past work (Bruce et al., 2021) has demonstrated that relatively accurate reconstructions of tibia-fibula surfaces can be obtained by fitting SSM to palpable landmarks via optimization procedures.

In this case study, we provide a similar application to that outlined by Bruce et al., (2021). We combine our SSM of the tibia-fibula with a set of measured palpable landmarks to reconstruct the entire surface. The ‘generateSurfaceFromLandmarks.m’ function provided with our dataset demonstrates an optimization procedure which fits the tibia-fibula SSM to a set of palpable landmarks on a new set of tibia-fibula surfaces (Nolte et al., 2020). We used the same set of palpable landmarks as Bruce et al., (2021) those being the tibial tuberosity, medial condyle, lateral and medial malleoli, lateral aspect of the fibula head, anterior border of the tibia at 25%, 50% and 75% of the distance between the medial condyle and malleolus, and the lateral fibula diaphysis at 25% of the distance from the lateral malleolus to the lateral point on the fibula head. These landmarks were digitized on the mean surface from the tibia-fibula SSM, as well as on 35 surfaces from the new dataset (Nolte et al., 2020). We performed nodal correspondence and registration of the new surfaces to the SSM mean using the coherent drift point algorithm and Procrustes analysis, respectively. The principal component scores from the SSM were then manipulated within an optimization that minimized the summed Euclidean distance between the landmarks on the SSM mean surface and new surface.

The mean and summed error in landmark positions (in mm) were 4.29 [4.01, 4.58 95% CIs] and 38.65 [36.09, 41.22 95% CIs], respectively; while the Jaccard index values for the entire reconstruction were 0.537 [0.517, 0.557 95% CIs] (see row 1 of Figure 8). Mean and peak error in surface node position (in mm) for the tibia surface were 6.63 [5.88, 7.39 95% CIs] and 13.39 [12.12, 14.65 95% CIs], respectively. The Jaccard index for the reconstructed tibia surfaces was 0.638 [0.620, 0.657 95% CIs] (see row 2 of Figure 8). Mean and peak error in surface node position (in mm) for the fibula, and the Jaccard index for the reconstructed fibula surfaces were 10.34 [8.85, 11.84 95% CIs], 20.82 [17.97, 23.67 95% CIs] and 0.271 [0.240, 0.303 95% CIs], respectively (see row 3 of Figure 8).

**Figure 8.**
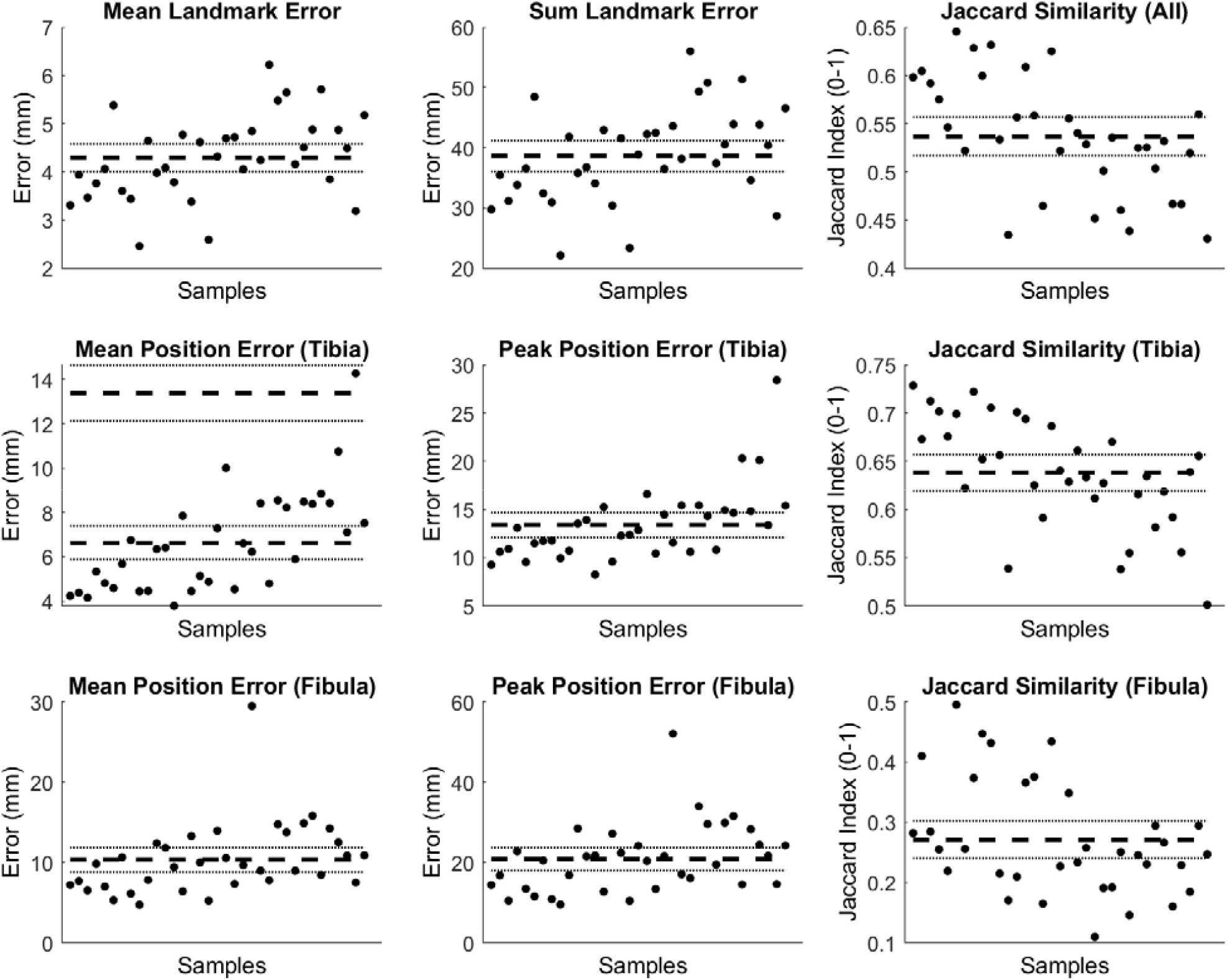
The mean and summed error, and Jaccard index for landmark positions (Row 1). Mean and peak error of the surface node position, and the Jaccard index for the tibia surfaces (Row 2) and fibula surface (Row 3) reconstructed using the palpable landmarks. Error variables are presented in mean (dashed line) and 95% confidence intervals (dotted lines).

This case study provides an example of how our statistical shape model can be used with minimal set of palpable landmarks on the tibia and fibula to estimate a 3D surface of the bones. Both the tibia and fibula can be reconstructed with reasonable accuracy. However, similar to Bruce et al., (2021), we noted better reconstruction performance and accuracy for the tibia over the fibula. The greater number and spread of landmarks on the tibia (i.e. 7 on the tibia vs. 2 on the fibula), and focus of the SSM components on the tibia are the likely reasons for this. Curvature of the fibula and relative position of the fibula head were common areas where larger discrepancies occurred (see Figure 9). Estimating additional landmarks at further points along the fibula or altering the code to give different weightings across landmarks in the optimisation may help address these issues. These may be particularly important for those who wish to use this code that have a specific interest in fibula anatomy and function. We also noted variation between samples with respect to reconstruction accuracy (see Figures 9 and 10). The variable performance was likely due to how well the original SSM captured the shape characteristics present in the individual samples.

**Figure 9.**
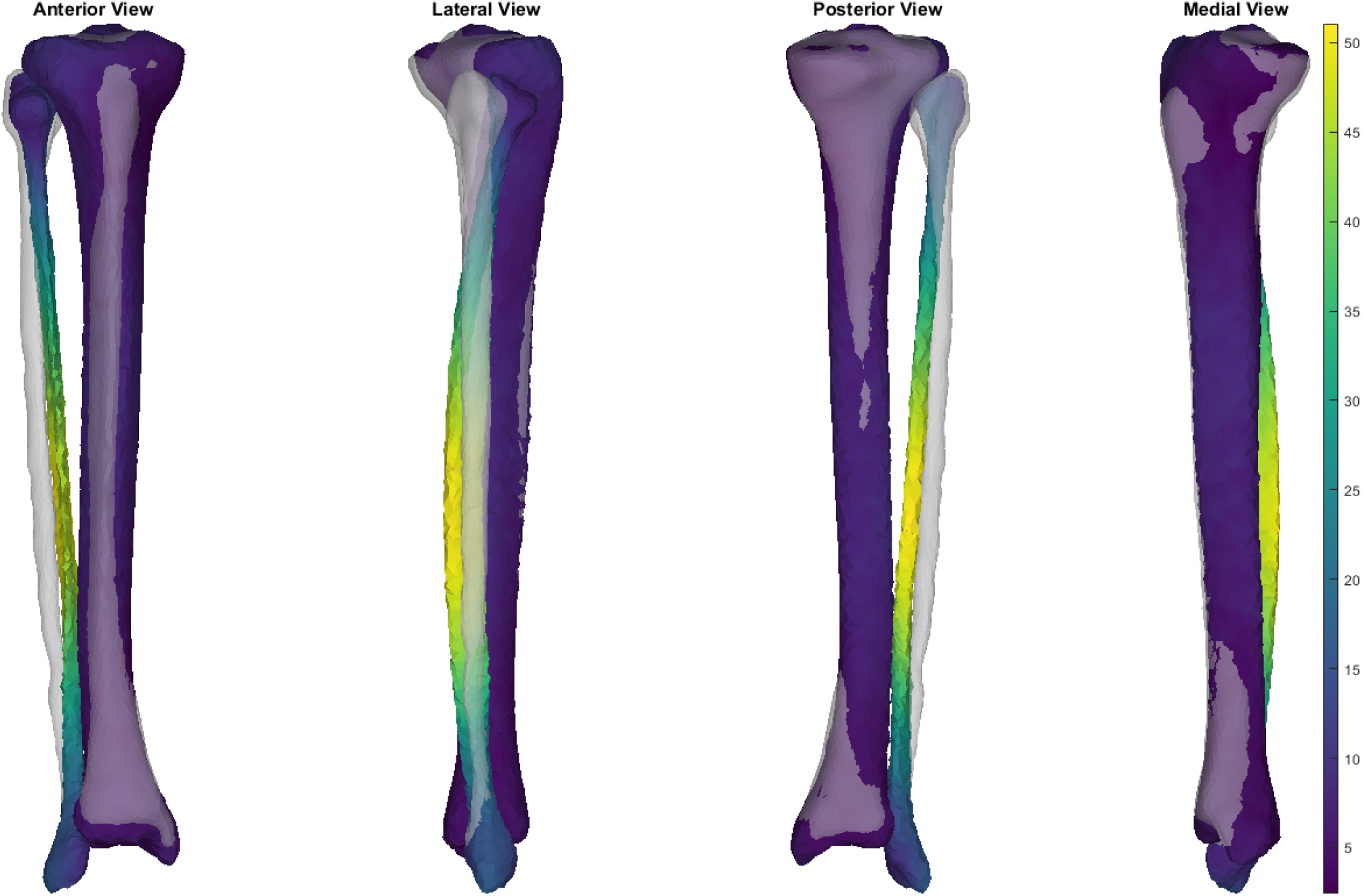
Example of a ‘poor’ reconstruction of the fibula from a minimized set of palpable landmarks. The transparent grey surface is the original segmentation, and the color map is the reconstructed surface. Warmer colours represent greater point distance (mm) from the original segmentation.

**Figure 10.**
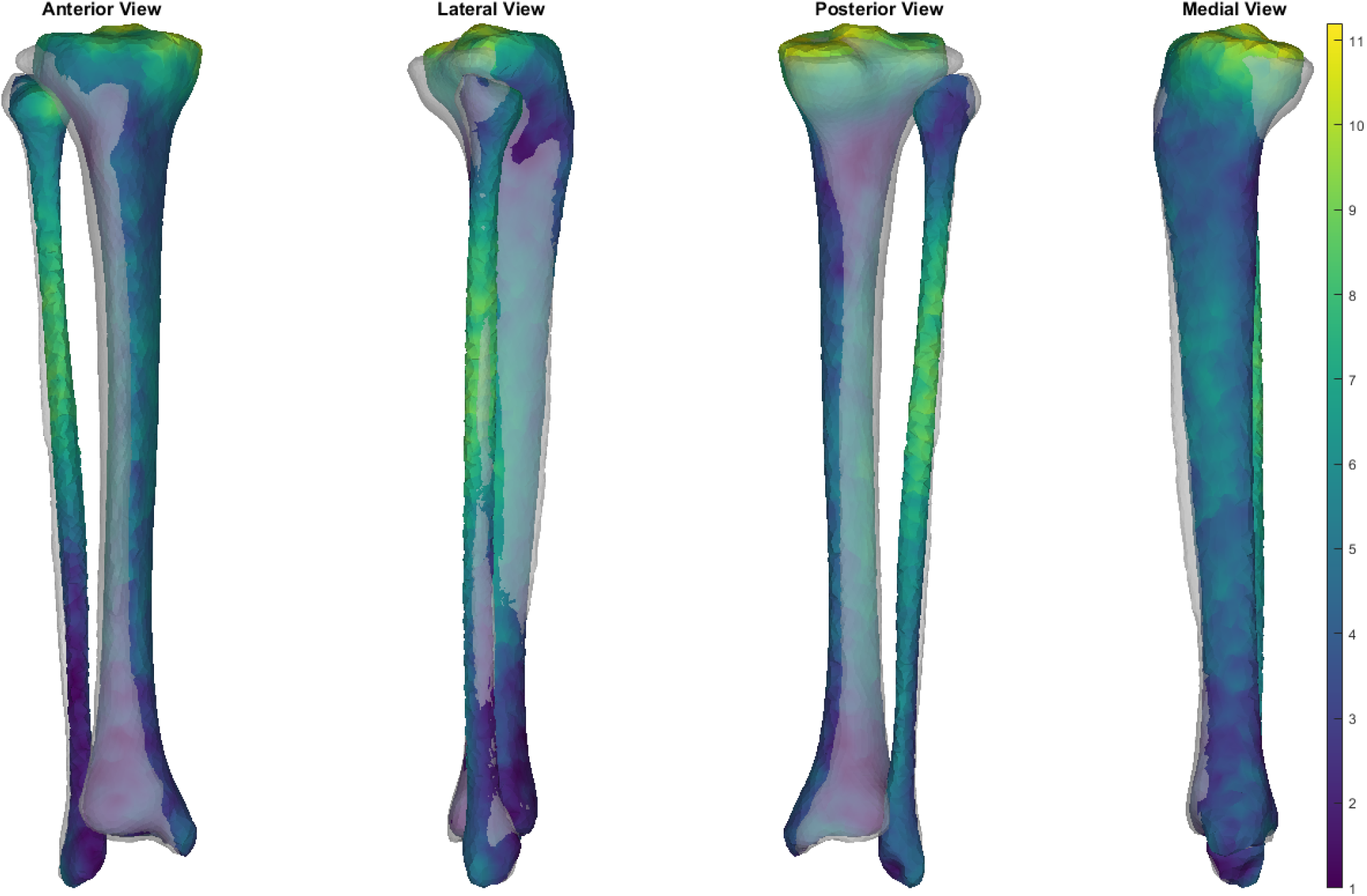
Example of a ‘good’ reconstruction of the tibia and fibula from a minimised set of palpable landmarks. The transparent grey surface is the original segmentation, and the color map is the reconstructed surface. Warmer colours represent greater point distance (mm) from the original segmentation.

### Limitations

The cadavers used were predicted to be from healthy adults (using meta-data provided by NMDID). Most participants meta-data included past medical history; however, we cannot be sure of the accuracy of all details. Hence, we cannot confirm that participants had no prior illness or injury that effected bone growth and formation.

The tibia-fibula set used to create the SSM was comprised of adults between the ages of 19 and 40. This may limit the applicability of the SSM outcomes to groups outside this age range, particularly geriatric and pediatric populations. For example, Bruce et al., 2021 demonstrated that surface reconstruction accuracy decreased when their SSM was applied to older populations, particularly in those whose bone may be affected by age-related degradation. Ethnicity is also known to effect bone geometry (Mahfouz et al., 2011; Sintini et al., 2018). While some included participants were born outside the United States, no data on ethnicity was available. The SSM should therefore not be used to examine ethnic factors related to bone geometry.

Lastly, due to only using the right tibia-fibula, researchers would be unable to use the code and SSM to reconstruct or perform analysis on a left tibia-fibula.

## Conclusions

Across all three of our SSM the largest source of geometric variation was a general size variation, with changes in both length and width observed. Important variations that could increase the risk of tibial stress injury were observed in the tibia and tibia-fibula SSM. These included general tibial thickness, midshaft thickness, tibial length, and medulla cavity diameter (indicative of cortical thickness). Further research is warranted to better understand the effect of these tibial and fibula shape characteristics on tibial stress, loading and injury risk. This SSM and associated code has been provided in an opensource dataset for use within the research community. The associated code includes three example applications, these include generation of a random sample, reconstruction of trabecular surfaces and reconstruction from palpable landmarks. In providing this dataset we hope to improve research skills in those individuals who may not have access to the knowledge or training. Further, this dataset could help to improve understanding of bone stress injuries, and potentially assist other research such as the implementation of medical devices (i.e. plates and braces). The SSM and all available code will be available for use at (https://simtk.org/projects/ssm_tibia.).

## Acknowledgements

The authors would like to acknowledge and thank the creators of the New Mexico Decedent Image database for providing the CT imagery used to conduct this research.

