## Supplementary figures and images for "Geometric variation of the human tibia-fibula: A public dataset of tibia-fibula surface meshes and statistical shape model"

### PC1_minus-3SD_animation.gif

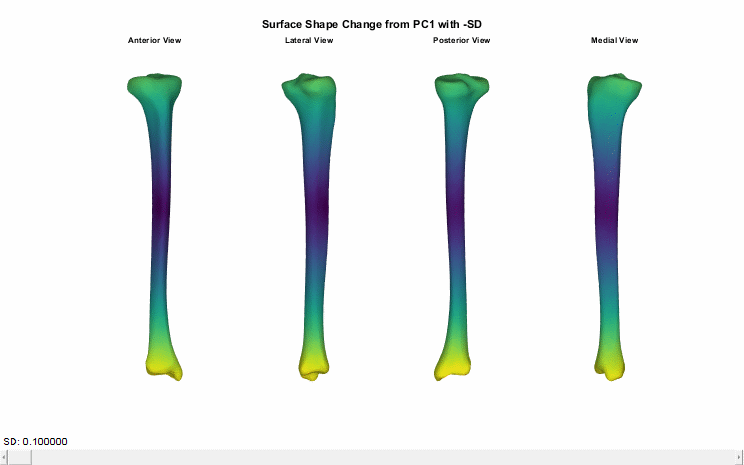

### PC1_minus-3SD_animation.gif

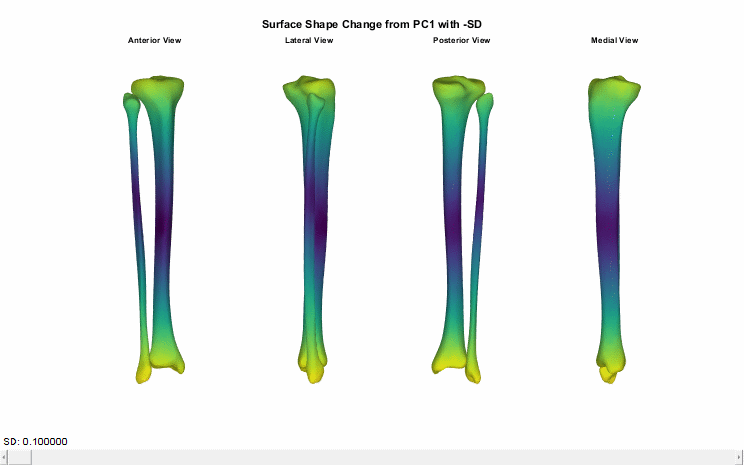

### PC1_minus-3SD_animation.gif

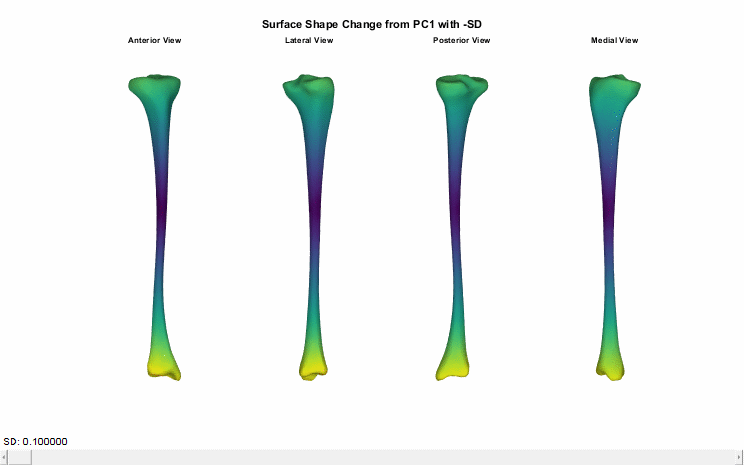

### PC1_plus-3SD_animation.gif

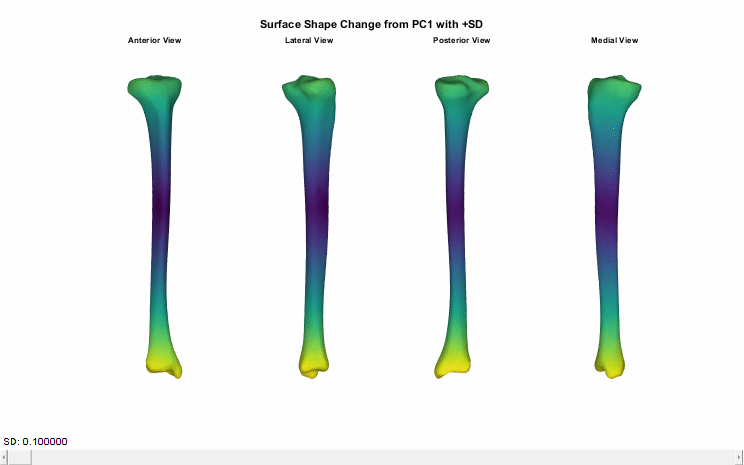

### PC1_plus-3SD_animation.gif

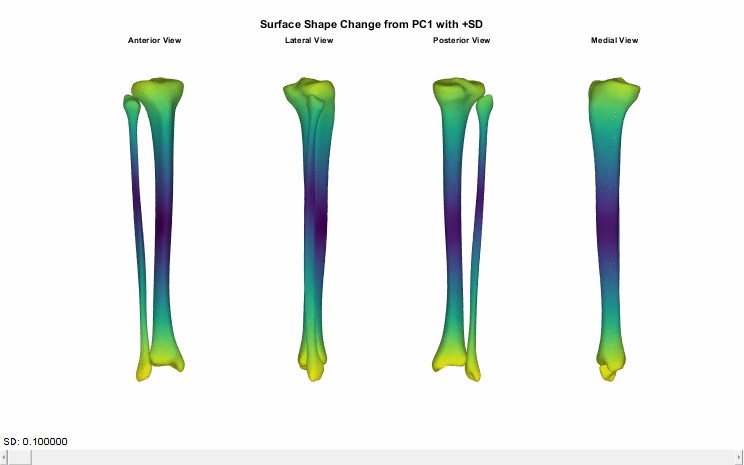

### PC1_plus-3SD_animation.gif

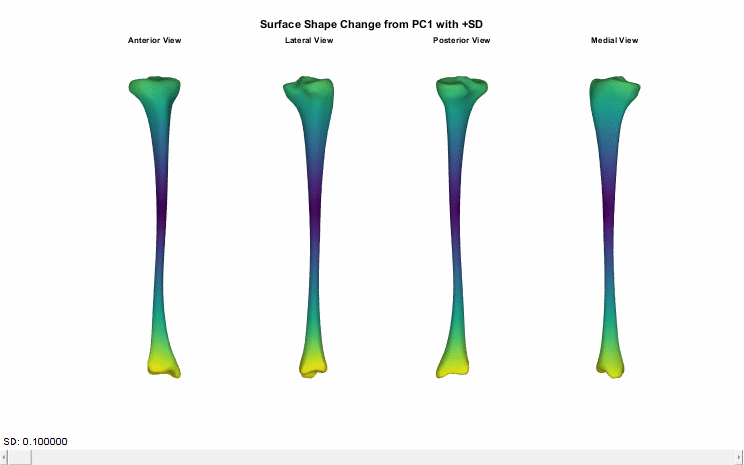

### PC2_minus-3SD_animation.gif

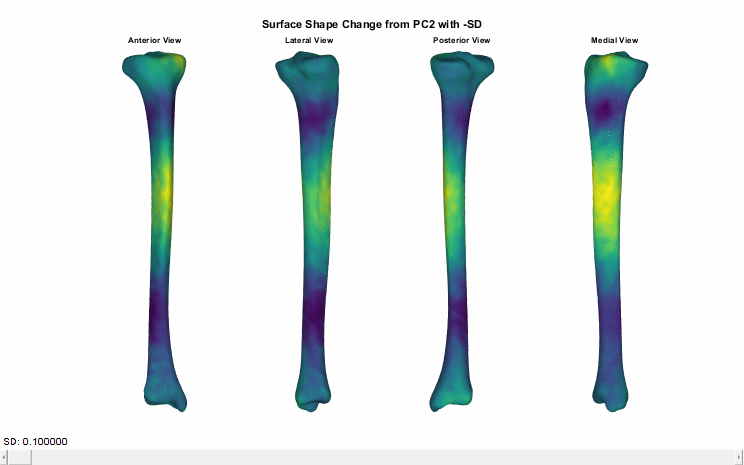

### PC2_minus-3SD_animation.gif

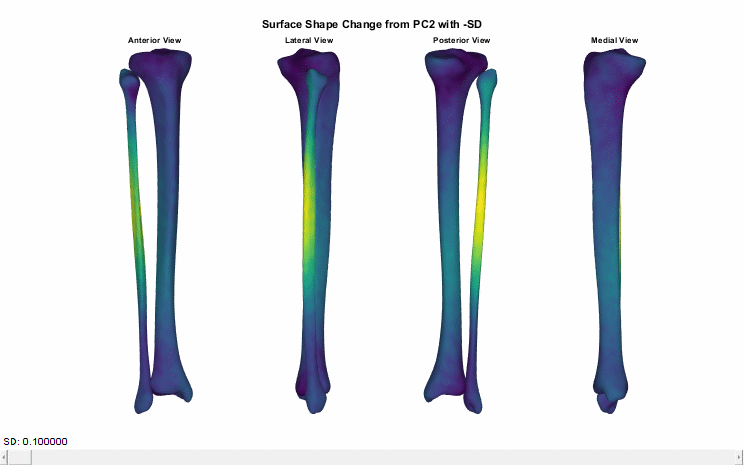

### PC2_minus-3SD_animation.gif

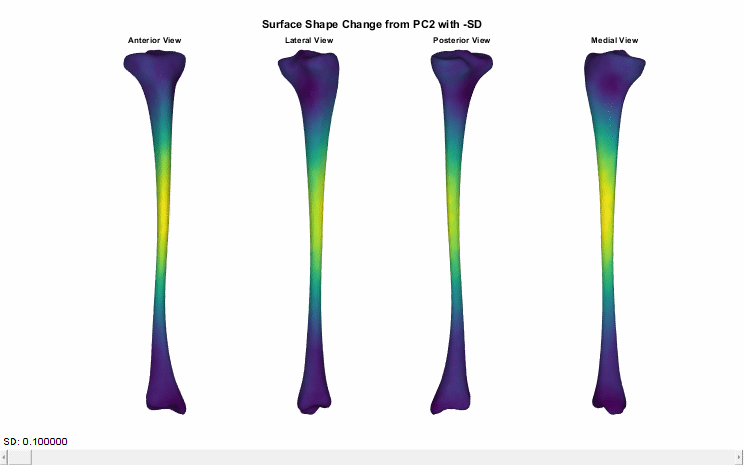

### PC2_plus-3SD_animation.gif

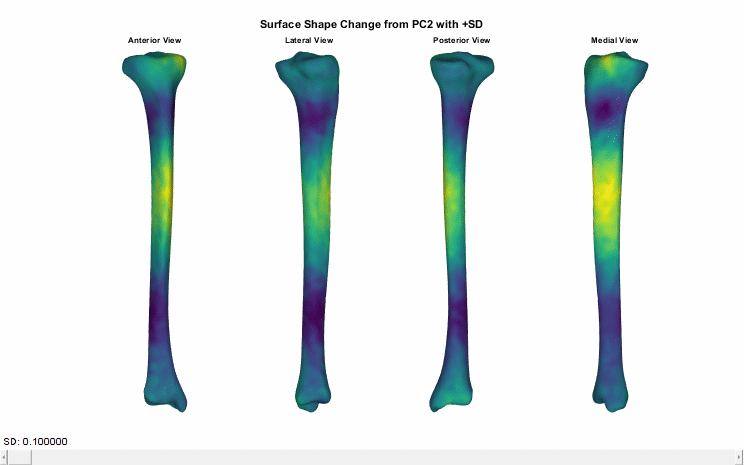

### PC2_plus-3SD_animation.gif

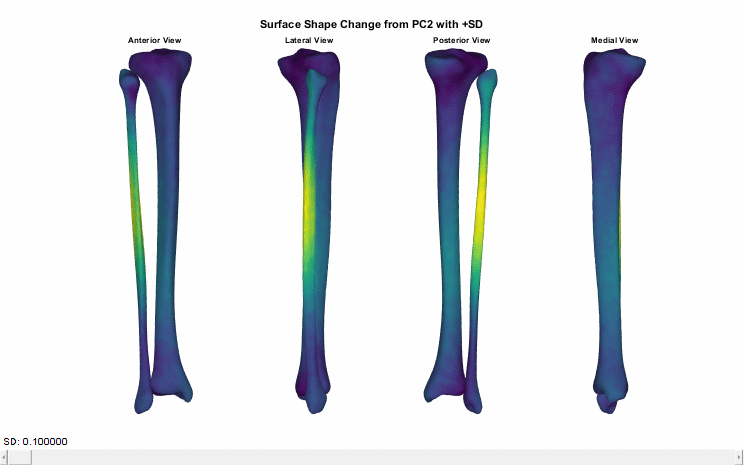

### PC2_plus-3SD_animation.gif

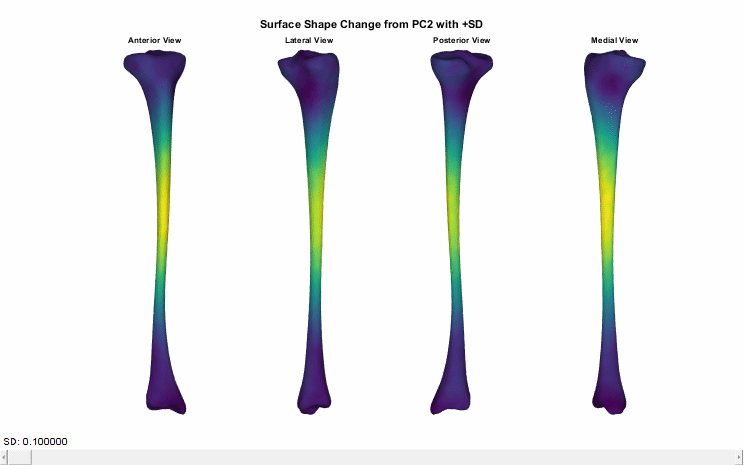

### PC3_minus-3SD_animation.gif

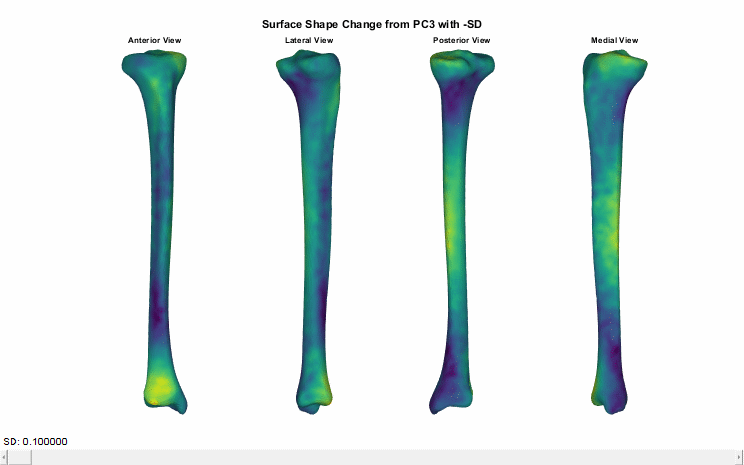

### PC3_minus-3SD_animation.gif

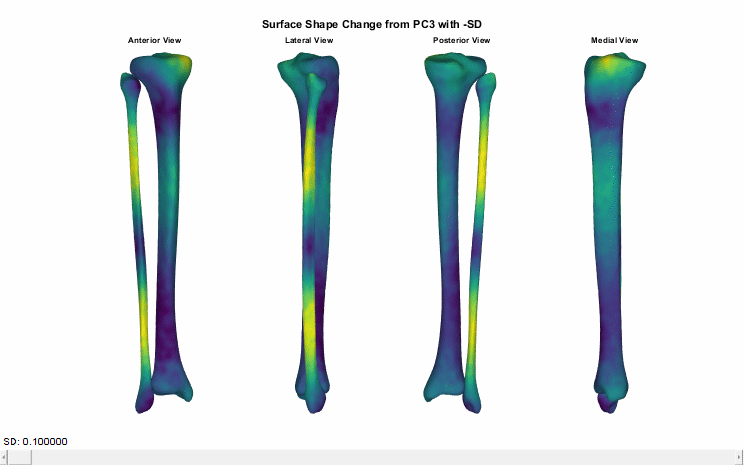

### PC3_minus-3SD_animation.gif

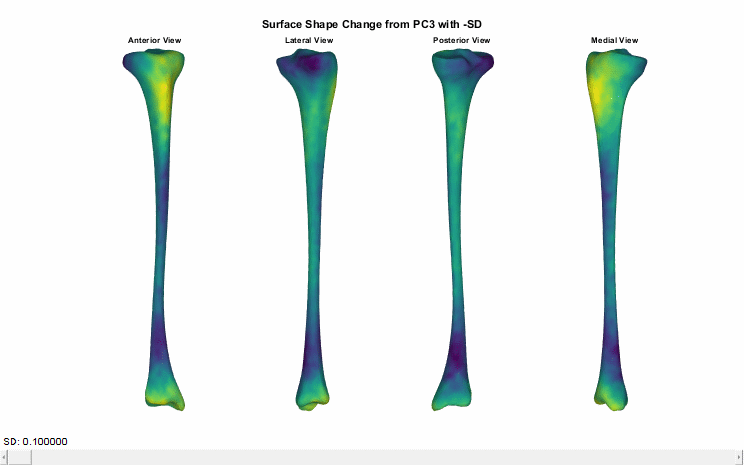

### PC3_plus-3SD_animation.gif

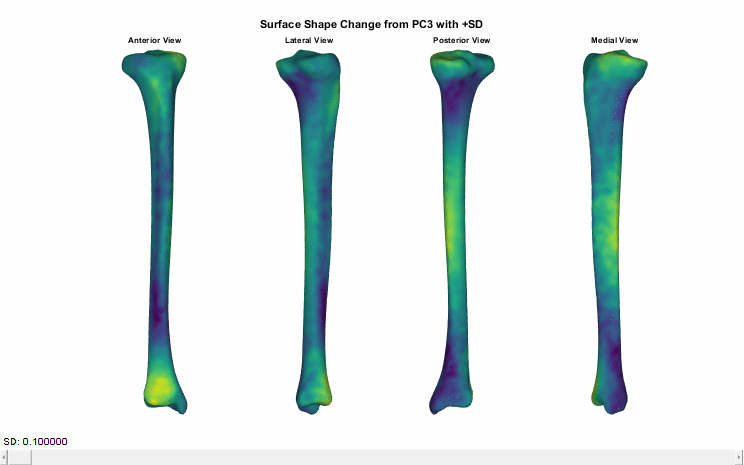

### PC3_plus-3SD_animation.gif

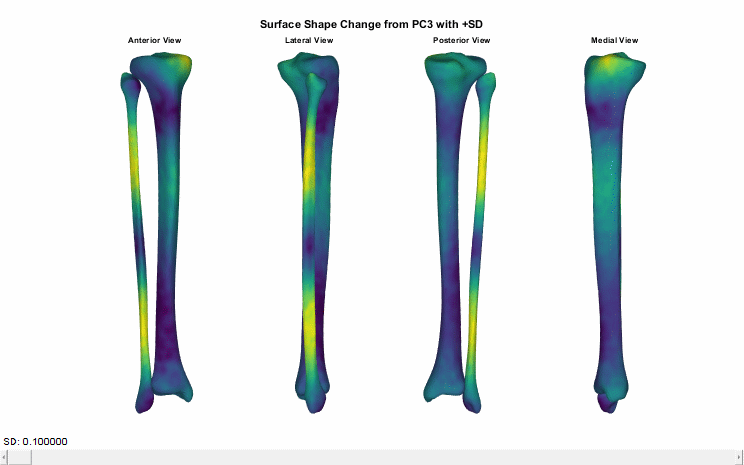

### PC3_plus-3SD_animation.gif

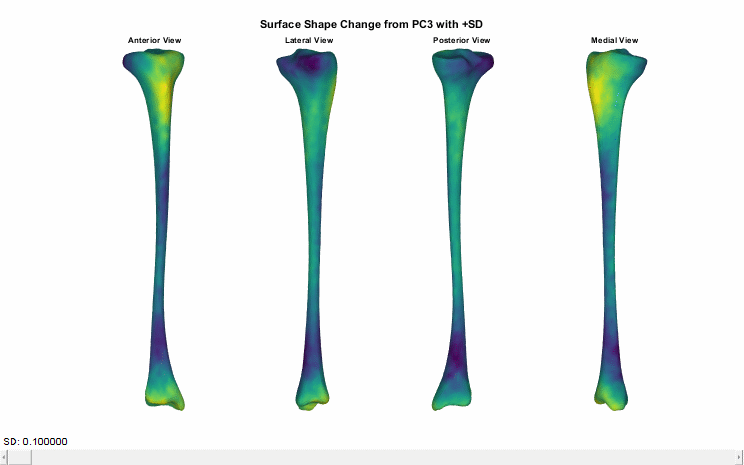

### PC4_minus-3SD_animation.gif

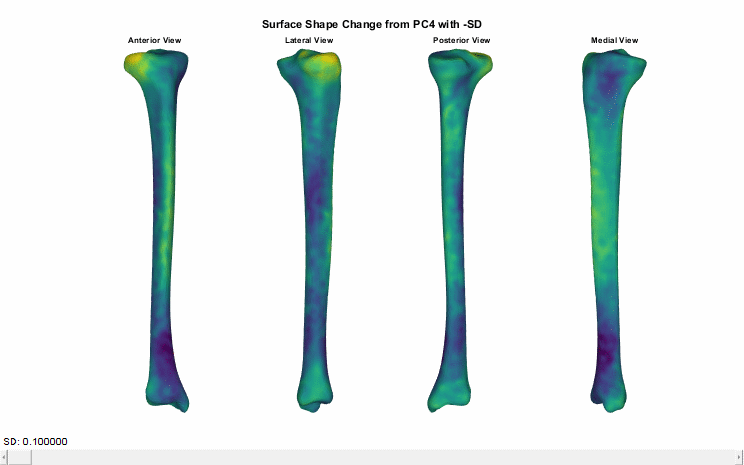

### PC4_minus-3SD_animation.gif

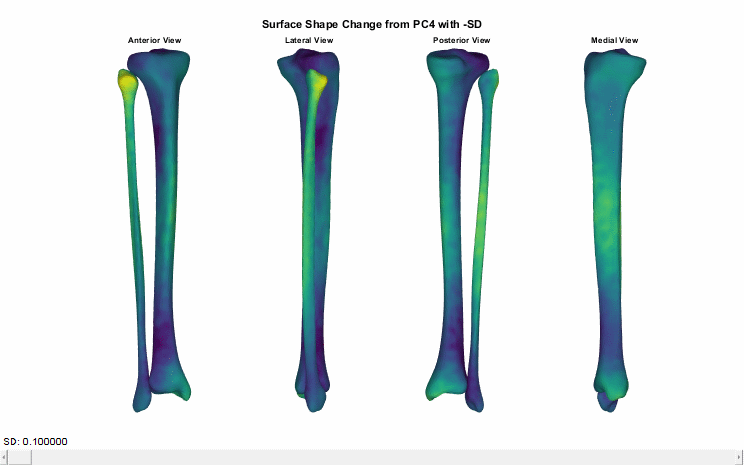

### PC4_minus-3SD_animation.gif

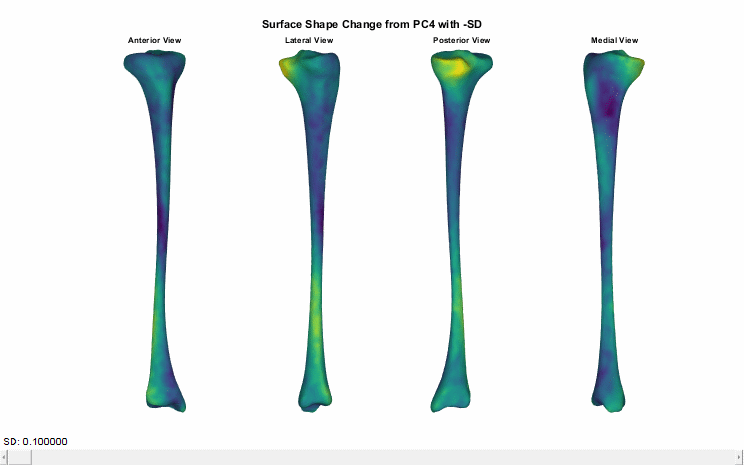

### PC4_plus-3SD_animation.gif

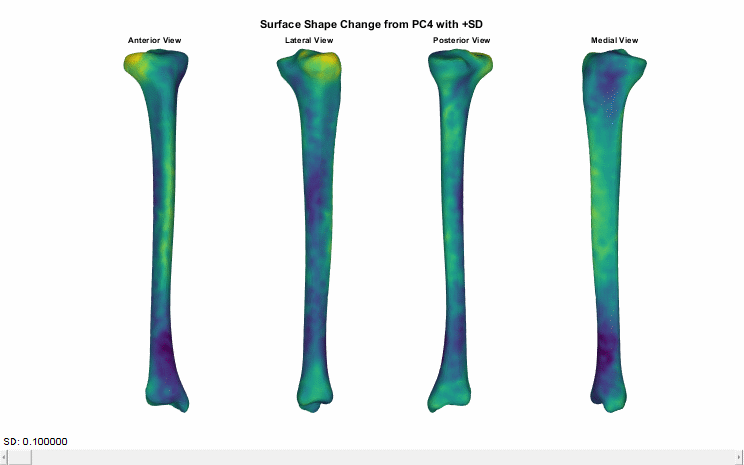

### PC4_plus-3SD_animation.gif

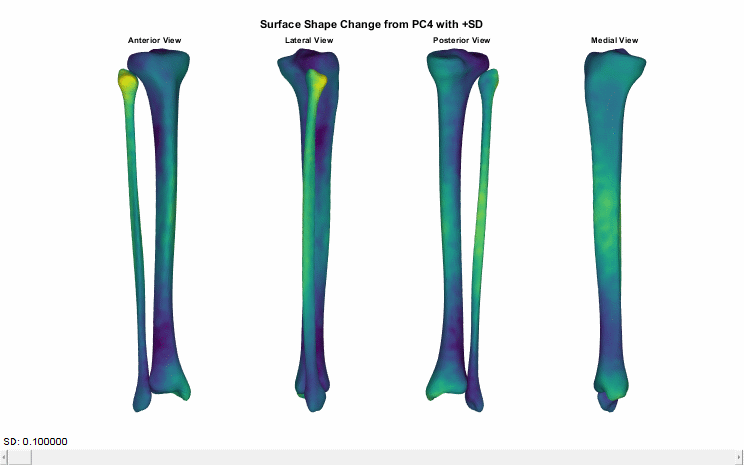

### PC4_plus-3SD_animation.gif

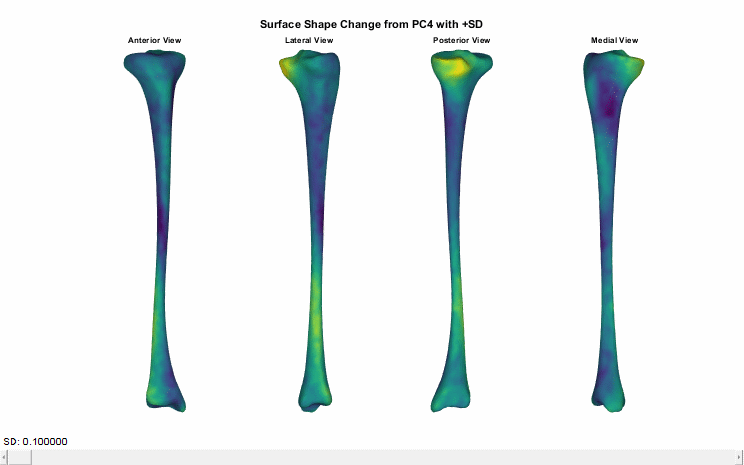

### PC5_minus-3SD_animation.gif

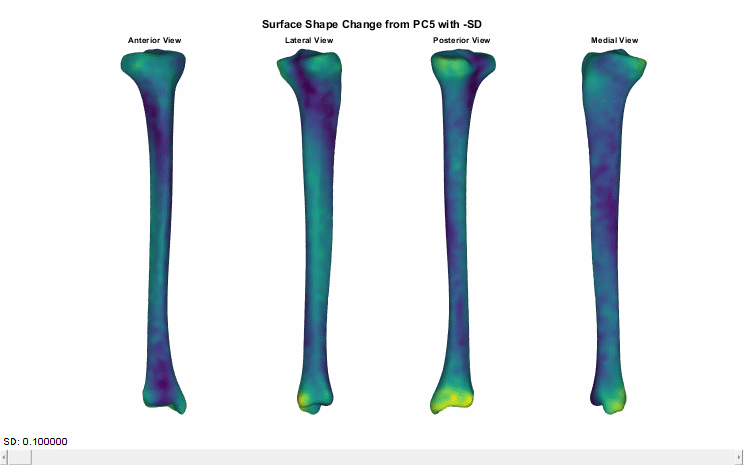

### PC5_minus-3SD_animation.gif

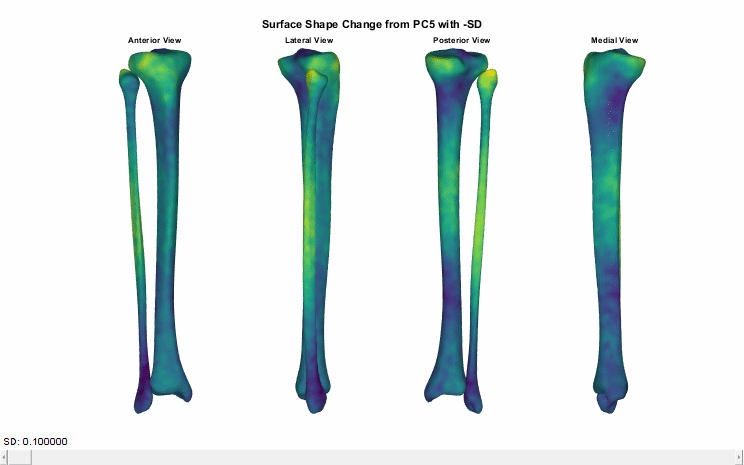

### PC5_plus-3SD_animation.gif

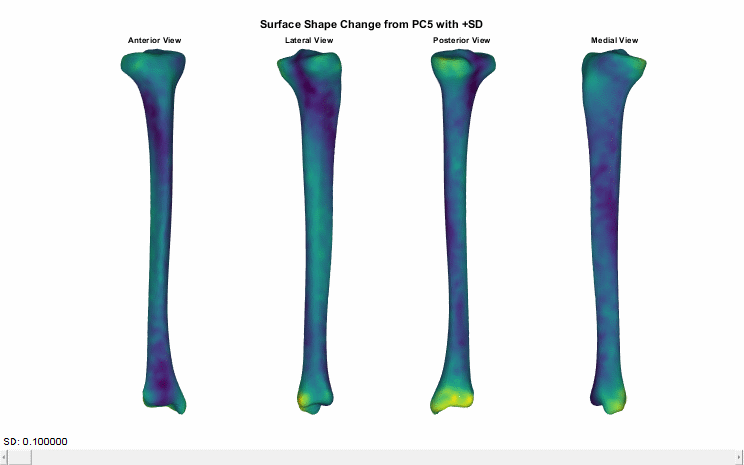

### PC5_plus-3SD_animation.gif

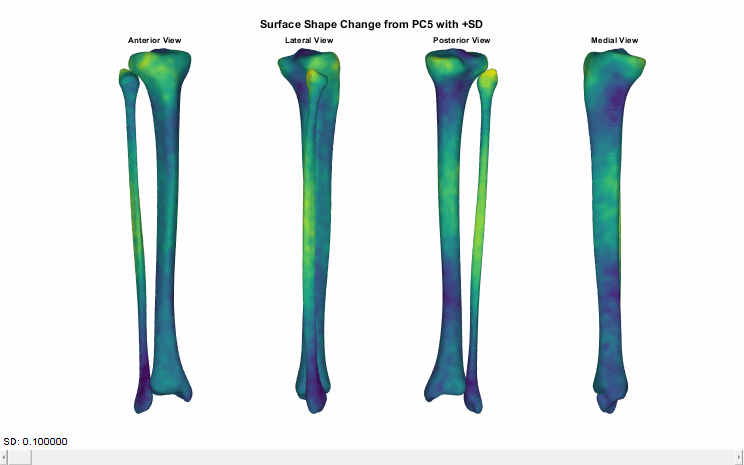

### PC6_minus-3SD_animation.gif

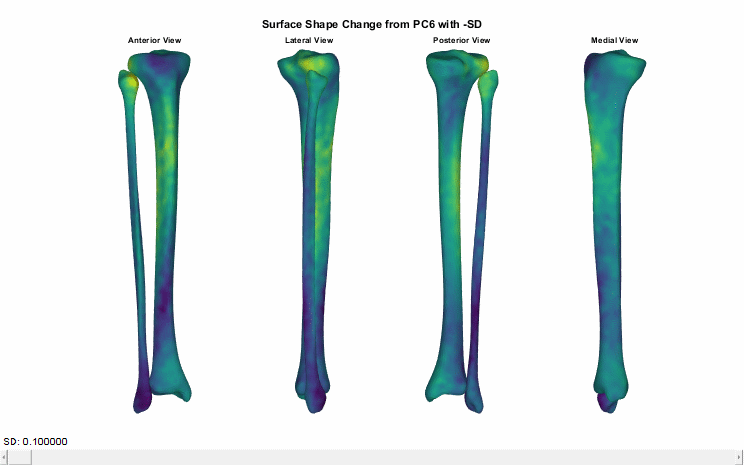

### PC6_plus-3SD_animation.gif

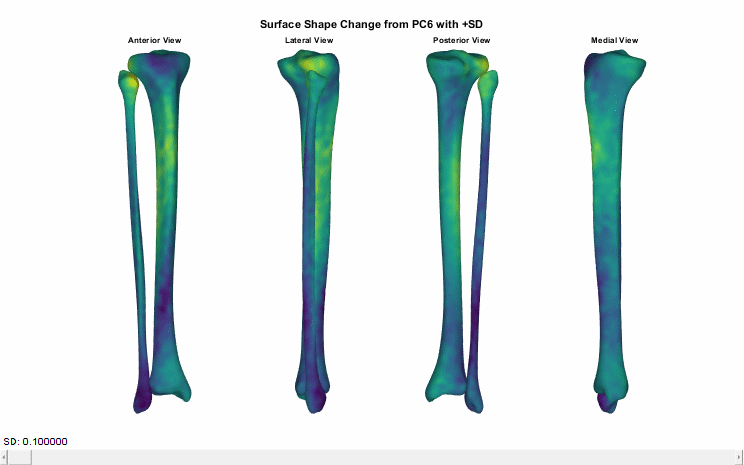

### PC7_minus-3SD_animation.gif

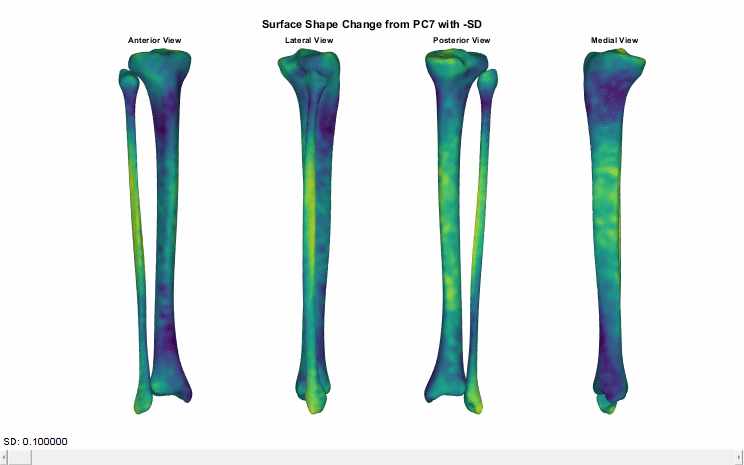

### PC7_plus-3SD_animation.gif

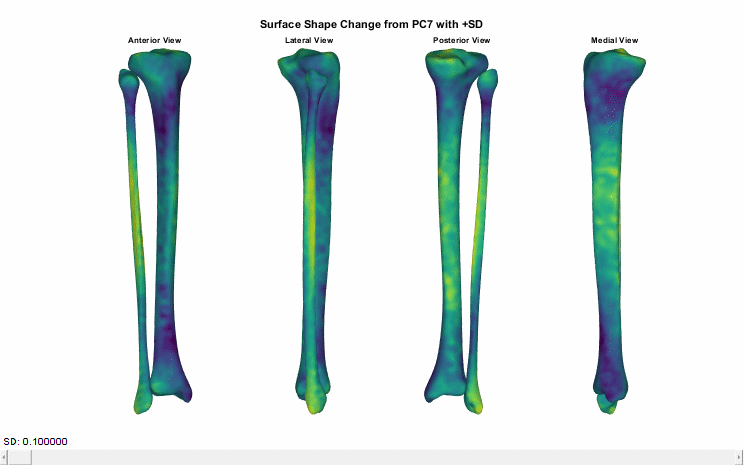
