## Supplementary File 4 for "Geometric variation of the human tibia-fibula: A public dataset of tibia-fibula surface meshes and statistical shape model"

### Supplementary Material 4

Below are the linear regression models for predicting each of the four retained trabecular principal shape model components from the five retained tibia shape model principal components.

#### Linear regression model summary for predicting trabecular shape model principal component (PC) 1 from the five retained tibia shape model PCs

| | Estimate | Standard Error | t-statistic | p-value |
|---|---|---|---|---|
| ***Intercept*** | *6.23E-14* | *8.018073* | *7.77E-15* | *1* |
| **Tibia Shape Model PC1** | 1.051309 | 0.015142 | 69.43087 | 3.53E-29 |
| **Tibia Shape Model PC2** | -0.00911 | 0.094809 | -0.0961 | 0.924239 |
| **Tibia Shape Model PC3** | -0.21684 | 0.126368 | -1.71596 | 0.099053 |
| **Tibia Shape Model PC4** | -0.13368 | 0.147469 | -0.9065 | 0.373683 |
| **Tibia Shape Model PC5** | 0.195048 | 0.148909 | 1.309849 | 0.202647 |

Number of observations: 30; Error degrees of freedom: 24

Root Mean Squared Error: 43.9

R-squared: 0.995; Adjusted R-Squared: 0.994

F-statistic vs. constant model: 965, p-value = 7.76e-27

**Linear regression model summary for predicting trabecular shape model principal component (PC) 2 from the five retained tibia shape model PCs**

| | Estimate | Standard Error | t-statistic | p-value |
|---|---|---|---|---|
| ***Intercept*** | 5.19E-15 | 23.77471 | 2.18E-16 | 1 |
| **Tibia Shape Model PC1** | 0.001674 | 0.044898 | 0.037276 | 0.970573 |
| **Tibia Shape Model PC2** | 1.248243 | 0.281123 | 4.440209 | 0.000172 |
| **Tibia Shape Model PC3** | 0.537295 | 0.374698 | 1.43394 | 0.164491 |
| **Tibia Shape Model PC4** | 0.706236 | 0.437266 | 1.615117 | 0.119357 |
| **Tibia Shape Model PC5** | -0.22721 | 0.441536 | -0.51459 | 0.611546 |

Number of observations: 30, Error degrees of freedom: 24

Root Mean Squared Error: 130

R-squared: 0.507; Adjusted R-Squared: 0.404

F-statistic vs. constant model: 4.93, p-value = 0.00304

**Linear regression model summary for predicting trabecular shape model principal component (PC) 3 from the five retained tibia shape model PCs**

| | Estimate | Standard Error | t-statistic | p-value |
|---|---|---|---|---|
| ***Intercept*** | -1.82E-14 | 8.329954208 | -2.18E-15 | 1 |
| **Tibia Shape Model PC1** | 0.002526837 | 0.015730782 | 0.160630106 | 0.873729155 |
| **Tibia Shape Model PC2** | -0.552013339 | 0.098497064 | -5.60436339 | 9.08E-06 |
| **Tibia Shape Model PC3** | 0.574078862 | 0.13128323 | 4.372827073 | 0.000204677 |
| **Tibia Shape Model PC4** | 0.163143328 | 0.153205064 | 1.064869033 | 0.297531822 |
| **Tibia Shape Model PC5** | 0.292330506 | 0.15470126 | 1.889645281 | 0.070944419 |

Number of observations: 30, Error degrees of freedom: 24

Root Mean Squared Error: 45.6

R-squared: 0.697; Adjusted R-Squared: 0.634

F-statistic vs. constant model: 11.1, p-value = 1.32e-05

**Linear regression model summary for predicting trabecular shape model principal component (PC) 4 from the five retained tibia shape model PCs**

| | Estimate | Standard Error | t-statistic | p-value |
|---|---|---|---|---|
| ***Intercept*** | -4.22E-14 | 9.716080904 | -4.34E-15 | 1 |
| **Tibia Shape Model PC1** | -0.00726552 | 0.018348426 | -0.3959751 | 0.695621756 |
| **Tibia Shape Model PC2** | -0.136737139 | 0.114887239 | -1.190185608 | 0.245613771 |
| **Tibia Shape Model PC3** | -0.466065694 | 0.153129111 | -3.043612613 | 0.005593592 |
| **Tibia Shape Model PC4** | -0.180973235 | 0.178698797 | -1.012727777 | 0.321291886 |
| **Tibia Shape Model PC5** | 0.477966417 | 0.180443964 | 2.648835719 | 0.014057495 |

Number of observations: 30, Error degrees of freedom: 24

Root Mean Squared Error: 53.2

R-squared: 0.44; Adjusted R-Squared: 0.324

F-statistic vs. constant model: 3.78, p-value = 0.0115
